# The Initial Detection of a Diversity of Viruses Associated with Ladybirds and Other Biocontrol Agents Prompts Interesting Ecological Insights

**DOI:** 10.1101/2025.10.06.680744

**Authors:** Gabriela B. Caldas-Garcia, Ícaro Santos Lopes, Eric Roberto Guimarães Rocha Aguiar

## Abstract

Predatory insects such as ladybirds, green lacewings, and an aphid-feeding gall midge have been commercialized and deployed worldwide to deal with the challenge of pest control. However, no information is available on their viruses which could interfere with the efficiency of biocontrol. To start filling this gap, this study employed an integrative bioinformatics approach starting from 21 publicly available RNA-seq libraries from 6 countries of 8 natural enemy species used to control aphids. A total of 41 putative novel viruses were identified and classified into 16 families and 1 genus. In addition, 13 known viruses were detected, including aphid and phytopathogenic viruses, and a virus from the invasive harlequin ladybird. The Aphid lethal paralysis virus was detected in several species at different trophic levels, including plants, aphid vectors, predatory insects, and insectivorous vertebrates. Thus, an ecosystem perspective on viruses across trophic levels is proposed because feeding may facilitate the spread of viruses between distant taxonomic species, such as invertebrate and vertebrate insectivorous predators. In conclusion, a simple assessment of the viral diversity in eight predatory insects of agricultural relevance, using a bioinformatics approach, has revealed a potential interspecies viral flux that deserves research attention. Further studies on viruses circulating in agricultural insect species are essential to improve biological control strategies based on trophic interactions, to conserve biodiversity, and to develop knowledge on the evolution of viral entities and their role as members of an agroecological system.

## 1. Introduction

Ladybirds (Coleoptera: Coccinellidae), lacewings (Neuroptera: Chrysopidae), and aphid midges (Diptera: Cecidomyiidae) have been recognized, commercialized, and applied as biological control agents, mainly to suppress soft-bodied insects, in several countries [1–10]. As a result, multiple biological control strategies have been successfully employed for over a hundred years in both open fields and greenhouses [11–13]. Aphids (Hemiptera: Aphididae) are critical pests of economically important and widely cultivated crops such as cereals, fruits, and legumes. The exponential reproductive rate and short generation time of aphids make them difficult to control [14], and because they feed on the phloem sap of host plants, they act as insect vectors of several viruses [15–17]. Although diverse aphid viruses have been identified over the past 40 years, knowledge on the effects of viral infection on these organisms remains limited [18]. Even more scarce is the knowledge about any viruses that infect essential aphidophagous insects, like ladybirds and lacewings.

As a voracious predator in both larval and adult stages, a ladybird can consume about 50 aphids a day, or up to 40,000 over a lifespan of about one year [19,20]. Ladybirds can help with pest control in several crops, such as cereals, fruits, potatoes, and sugar beet [20,21]. However, negative effects on the diversity and abundance of coccinellids have been reported around the world [22–24], mostly due to the occurrence of invasive species (such as *Harmonia axyridis* [25]), overuse of agrochemicals [26–28], urbanization [29], and climate change [30]. For instance, the two-spotted lady beetle (*Adalia bipunctata*) was abundant in Europe, but over the last decades it has been declining in that continent and in North America [28,31]. Moreover, ladybirds are more vulnerable insects than well-studied honeybees because they go through larval development and adult life without the protection of a colony or a beekeeper [28,32].

A lacewing in its larval stage can eat between 500 to 1500 aphids for about 2 to 3 weeks [19,20]. This predator is found controlling harmful insects in cultivars of fruits, vegetables, nuts and others [33,34]. The diversity of lacewings has decreased since the Cretaceous (130–100 million years ago) accompanied by a great loss of ecological roles as well, since several morphologies of lacewing larvae have been lost through time [35]. Currently, they are also negatively affected by pesticide exposure [36–38]. Another biocontrol agent, the aphidophagous midge *Aphidoletes aphidimyza*, preys on about 85 aphid species and is released in a wide range of crops, including strawberries and ornamentals [2,20]. Each larva eats up to 80 aphids per day, while the adult feeds on honeydew and nectar [2,20]. More details on the predator species included in this work can be found in **Table 1** and its references [2,3,9,34,39–48].

**Table 1.** Information on the beneficial insects selected for virus analysis.

| Ladybirds (Coccinellidae) |  |  |  |  |  |  |  |  |
| --- | --- | --- | --- | --- | --- | --- | --- | --- |
| Species | Libraries* | Sample* | Collection* | Geographic location* | Geographic distribution | Ecosystem service provided | Consumed preys | References |
| *Coccinella septempunctata* | SRR12728770<br>Total RNA-seq | adult | *Vicia faba* | Beijing, China | Native to Europe and Asia, and introduced in the Nearctic region; Widespread | Biocontrol - polyphagous predator at the larval and adult stages | Aphids, whiteflies, thrips, and scale insects | Bauer, T. 2013 (On-line). Kundoo & Khan, 2017. |
|  | SRR9649799<br>poly (A) | 4th instar larva fed mealybugs | laboratory | Guangzhou, China |  |  |  |  |
|  | ERR1145729<br>poly (A) | adult whole beetle | field | Germany |  |  |  |  |
|  | ERR1145726<br>poly (A) | adult whole beetle | field | Germany |  |  |  |  |
| *Adalia bipunctata* | ERR1145720<br>poly (A) | adult whole beetle | laboratory | Germany | Native to Europe, Central Asia and North America | Biocontrol - both adult and larval forms are voracious aphidophagous hunters, polyphagous predator | Several aphid species, coccids, diaspid, and pollen | Riddick, 2017. Omkar & Pervez, 2005. |
|  | ERR1145719<br>poly (A) | adult whole beetle | laboratory | Germany |  |  |  |  |
| *Cryptolaemus montrouzieri* | SRR8325176<br>Total RNA-seq | 4th instar larva fed pollen | laboratory | Guangzhou, China | Native to Australia, introduced to at least 64 countries or regions for biocontrol purposes | Biocontrol - effective predator of mealybugs | At least 21 mealybug species, aphids, coccids, and whiteflies | Xie et al., 2017. Li et al., 2021. Kairo et al., 2013. |
|  | SRR24179305 | gut |  | Guangzhou, China |  |  |  |  |
| *Novius pumilus* | SRR15420501<br>Total RNA-seq | adult female whole body under starvation | lab-reared | Guangzhou, China | Native to China. *Novius* spp. are distributed worldwide | Biocontrol - predator of scale pests | Several coccid species, such as *Icerya aegyptiaca* , *I. purchasi*, and *I. seychellarum* | Tang et al., 2022. Tang et al., 2024. |
|  | SRR15420505<br>Total RNA-seq | adult female whole body fed *Icerya aegyptiaca* | lab-reared | Guangzhou, China |  |  |  |  |
|  | SRR15420491<br>Total RNA-seq | adult female whole body fed *Icerya aegyptiaca* | lab-reared | Guangzhou, China |  |  |  |  |
|  | SRR15420492<br>Total RNA-seq | adult female whole body fed *Icerya aegyptiaca* | lab-reared | Guangzhou, China |  |  |  |  |
| *Propylea japonica* | SRR18494825 | adult female whole body |  | Shaanxi, China | East of Europe and Asia | Biocontrol - larvae and adults are dominant natural predators in China | Aphids, whiteflies, scale insects, planthoppers, and lepidopteran larva | Huangfu et al., 2023. |
|  | SRR1299012<br>poly (A) | body of all stages of ladybird | laboratory | Guangzhou, China |  |  |  |  |
| Lacewings (Chrysopidae) |  |  |  |  |  |  |  |  |
| Species | Libraries* | Sample* | Collection* | Geographic location* | Geographic distribution | Ecosystem service provided | Consumed preys | References |
| *Chrysoperla carnea* | SRR10012087<br>poly (A) | head and body | laboratory | Belgium | Worldwide | Biocontrol - predatory, polyphagous at the larval stage | Insect and mite eggs, aphids, and other several soft-bodied insects (such as caterpillars, coccids, mealybugs, psyllids), mites and other arthropods | Amarasekare & Shearer, 2013. Tauber et al., 2000. |
|  | ERR12321221<br>poly (A) | adult whole body | field - Wytham Woods | Berkshire, United Kingdom |  |  |  |  |
|  | SRR1811846<br>poly (A) | adult whole body |  | Vienna, Austria |  |  |  |  |
| *Chrysopa pallens* | SRR5581488<br>poly (A) | whole body | laboratory | Beijing, China | Widespread - Asia, North America and Europe | Biocontrol - predator at both the adult and larval stages | Mites and several insects, such as aphids, thrips, planthoppers, caterpillars, and whiteflies | Wang et al., 2021. Sarkar et al., 2019. |
|  | SRR1811843<br>poly (A) | whole body |  | Vienna, Austria |  |  |  |  |
| Aphid midge (Cecidomyiidae) |  |  |  |  |  |  |  |  |
| Species | Libraries* | Sample* | Collection* | Geographic location* | Geographic distribution | Ecosystem service provided | Consumed preys | References |
| *Aphidoletes aphidimyza* | SRR1695317<br>poly (A) | adult whole body | laboratory | North Carolina, USA | Holarctic region, Argentina, Chile, Brazil and New Zealand | Biocontrol only at the larval stage | 85 aphid species | Boulanger et al., 2019 |
|  | SRR27213331<br>poly (A) | pupa whole body | laboratory | Jinan, China |  |  |  |  |

\* Information retrieved from the SRA and BioSample (NCBI) public databases

Therefore, natural enemies play a prominent role in sustainable agriculture and global food security. However, anthropic activities have contributed to accelerating the rates of environmental degradation and extinction of species [24,30,49,50]. For instance, pesticide usage can increase insect susceptibility to parasites and pathogens [51]. Nevertheless, there is a shortage of studies on viruses of predatory insects, such as ladybirds, lacewings, and aphid midges. Currently, there is no virus described for *A. aphidimyza*, despite its use in more than 20 countries [52]. On lacewing viruses, Shi et al. (2016) discovered the flavi-like Shuangao lacewing virus 2 in Chrysopidae sp. [53]. Finally, regarding ladybird viruses, there are two picorna-like viruses described in *Harmonia axyridis* (Harmonia axyridis virus 1) [54], and in *Cheilomenes sexmaculata* (Cheilomenes sexmaculata picorna-like virus 1) [55]. In addition, Dheilly et al. (2015) detected in *Coleomegilla maculata* the iflavirus Dinocampus coccinellae paralysis virus, which is transmitted by the endoparasitoid wasp *Dinocampus coccinellae* [56]. Given the paucity of current knowledge about viruses associated with predatory insects, and the lack of information regarding virus sharing among aphidophagous organisms, this study aims to (a) investigate what exogenous viruses are present in eight important beneficial insects using public next-generation sequencing RNA libraries; (b) analyze common viruses among generalist predatory species that occupy the same ecological niche (predators of soft-bodied insects, especially aphids); and (c) track other hosts and possible transmission/dispersal routes that could explain the presence and/or prevalence of the virus in its different hosts.

## 2. Materials and Methods

### 2.1 Data processing of public RNA-seq libraries

Twenty-one paired-end RNA-seq libraries of long RNAs from eight species of beneficial predatory insects were selected for virus investigation and retrieved from the NCBI Sequence Read Archive (SRA) repository (https://www.ncbi.nlm.nih.gov/sra), accessed until January 2024. The accession numbers grouped by species are available in **Table 1**, and see **Supplementary Table 1** for more details on the libraries analyzed. The following tools implemented on the Galaxy Australia web-based platform were utilized [57]: (a) FastQC version 0.74 [58] used for raw reads’ quality assessment; (b) Trimmomatic v. 0.36.6 [59] used for quality filtering of bases with quality scores below 20, through sliding window trimming operation; (c) Bowtie2 v. 2.5.0 [60] used for reads’ mapping against each host’s genome, aiming to select only unmapped reads, which may be putative viral reads; (d) rnaviralSPAdes v. 3.15.5 [61], SPAdes v. 3.15.5 [62], and Trinity v. 2.15.1 [63] used to assemble possible viral genomes; (e) DIAMOND BLASTx v. 2.0.15 [64], setting E-values ≤ 10^-5^ and standard parameters, used for identification of viral sequences assembled, and queried against the complete viral RefSeq (protein) database downloaded from the NCBI (https://ftp.ncbi.nlm.nih.gov/refseq/release/viral/), accessed on 22 December 2023.

### 2.2 Manual inspection and improvement of putative viral genomes

To identify the most accurate viral sequences, the DIAMOND hits exhibiting similarity to retroviruses, transposons, and DNA viruses were excluded from further analysis, as they predominantly represent false positives in virome studies based on the RNA-seq technique. In addition, putative DNA viruses can be endogenous viral elements (EVEs), and these elements deserve a study focused on them. Thus, the remaining sequences were subjected to a second round of similarity analysis to confirm their viral origin through the online NCBI BLAST (https://blast.ncbi.nlm.nih.gov/Blast.cgi, accessed until November 2024). In the blast phase, BLASTn (nucleotide collection database) and BLASTx (non-redundant protein sequences database) were employed. Contigs exhibiting high identity in sequence similarity searches at the nucleotide level (greater than 91%), substantial coverage (over 80%, contingent upon genome completeness), and E-value = 0.0 were assigned to known viral species. Further analysis was performed on the sequences that showed similarity to RNA viruses (lower than 91% id). For each one, the open reading frames (ORFs) were predicted using ORFfinder (https://www.ncbi.nlm.nih.gov/orffinder, accessed until November 2024); further, conserved domains were verified using profile hidden Markov models with phmmer search (https://www.ebi.ac.uk/Tools/hmmer/search/phmmer), NCBI Conserved Domains search tools (https://www.ncbi.nlm.nih.gov/Structure/cdd/wrpsb.cgi), and InterPro classification of protein families (https://www.ebi.ac.uk/interpro/search/sequence/), all accessed until November 2024. The viral sequences found incomplete, considering length and ORF structure, in comparison with their closest relative by BLAST searches were subjected to a new round of assemblage. Twelve libraries containing partial viral sequences were reassembled using an integrative pipeline that includes several assemblers (such as metaSPAdes [65], metaviralSPAdes v. 3.15.5, rnaviralSPAdes [61], MEGAHIT v. 1.2.9 [66], and Trinity [63]) followed by transcript unification with Cap3 [67], as previously utilized in earlier works [68,69].

### 2.3 Inference of phylogenetic trees

Complete or nearly complete viral sequences (length ≥ 90% of related viruses’ genome length) and partial sequences of viruses (length ≥ 1000 nt) containing the RNA-dependent RNA polymerase (RdRp) conserved domain were selected for the phylogenetic analysis. The trees of putative novel viruses were constructed using the longest ORF of each new virus translated by Expasy – Translate tool (https://web.expasy.org/translate/, accessed in November, 2024), followed by protein sequences of related viruses obtained by online BLASTx, in conjunction with sequences acknowledged by the International Committee on Taxonomy of Viruses (ICTV), when feasible. The ICTV viruses were picked up among members of the same taxonomic family and/or closely related genera, from the Virus Metadata Resource (VMR_MSL39.v2_20240920, accessed on 23 September 2024). For outgroups, ICTV members of other families or distant genera were selected. More information on details of phylogenetic trees’ construction is available in **Supplementary Table 2**. ORFs (aa) containing the RdRp domain were aligned by MAFFT (https://www.ebi.ac.uk/jdispatcher/msa/mafft?stype=protein, accessed in November 2024), and AliView v. 1.28 [70] was employed for minimal alignment fixing and end trimming. Next, the maximum likelihood trees were built using IQ-TREE v. 1.6.12 [71] setting up 1000 ultrafast bootstrap replicates performed by UFBoot [72]. The best-fit model was estimated by ModelFinder [73], and chosen according to the Bayesian Information Criterion [74]. For tree visualization, rooting, and editing, FigTree v. 1.4.4 [75] and iTol v. 7.0 [76] were utilized. The animal shadows used for illustration are available in PhyloPic (https://www.phylopic.org/). Finally, the naming format utilized for new virus species followed the Latinized binomial nomenclature recommended by the ICTV [77].

### 2.4 Estimation of viral transcript abundance from RNA-seq reads

Reference transcriptomes available for *Coccinella septempunctata* (https://ftp.ncbi.nlm.nih.gov/genomes/all/GCF/907/165/205/GCF_907165205.1_icCocSept1.1/GCF_907165205.1_icCocSept1.1_rna_from_genomic.fna.gz, accessed in November 2024), and *Chrysoperla carnea* (https://ftp.ncbi.nlm.nih.gov/genomes/all/GCF/905/475/395/GCF_905475395.1_inChrCarn1.1/GCF_905475395.1_inChrCarn1.1_rna_from_genomic.fna.gz, accessed in November 2024) were downloaded and added to the respective viral sequences found in each of the two species. Since there is no transcriptome of reference for the other six host species included in this study, one full library of each predator was assembled with SPAdes v. 3.15.5 [62] and evaluated with TransDecoder v. 5.5.0 (https://github.com/TransDecoder/TransDecoder, accessed in November 2024) for identifying the most probable coding sequences. The resulting host transcriptomes added to the viral genomes were utilized for estimating the viral abundance, in comparison to selected host mRNAs, using Salmon quant v. 1.10.1 [78]. Two host’s marker genes, one polyadenylated - *CAM* (calmodulin), and one not polyadenylated – *H3-3A* (histone H3) were searched in the reference transcriptomes for comparison with the viral amounts. Afterward, the selected *C. septempunctata* markers (GenBank: XM_044906086.1 and XM_044902344.1, accessed in November 2024) were identified through BlastN for each Coccinellidae species evaluated. Similarly, the selected markers of *Chrysoperla carnea* (GenBank: XM_044884488.1 and XM_044878447.1, accessed in November 2024) were used for BlastN searches in the *Chrysopa pallens* transcriptome. For *Aphidoletes aphidimyza*, sequences of *Drosophila melanogaster* were used as references for BlastN searches (GenBank: NM_078986.3 and NM_001273153.1, accessed in November 2024). Finally, for visualization of transcript abundance, a heatmap was built using R’s ComplexHeatmap package [79], with the transcripts per million (TPM) values normalized using log10 TPM+1.

## 3. Results

### 3.1 The remarkable viral diversity detected in the predatory ladybirds, lacewings, and aphid midges

The data analysis of 21 RNA-seq libraries from 6 countries (Austria, Belgium, China, Germany, United Kingdom, and United States of America) unveiled sequences of 13 known viruses (**Table 2**), 41 putative new viruses, and 4 fragments (< 1000nt length) of viral origin (**Table 3**). New viruses have been classified into 16 families recognized by the ICTV and 1 unclassified genus (*Negevirus*). In detail, according to the Baltimore classification system [80,81], there were viruses from 9 families (*Picornaviridae, Dicistroviridae, Solinviviridae, Iflaviridae, Polycipiviridae, Secoviridae, Tymoviridae, Narnaviridae,* and *Solemoviridae*), and 1 genus (*Negevirus*) classified as single-stranded positive sense RNA viruses (ssRNA+); 3 families (*Lispiviridae, Phasmaviridae*, and *Rhabdoviridae*) of single-stranded negative sense RNA viruses (ssRNA-); and 4 families (*Spinareoviridae, Partitiviridae, Amalgaviridae*, and *Orthototiviridae*) of double-stranded RNA viruses (dsRNA). **Figure 1** shows the characterization of the ORFs and domains for each virus found in this study.

**Figure 1.**
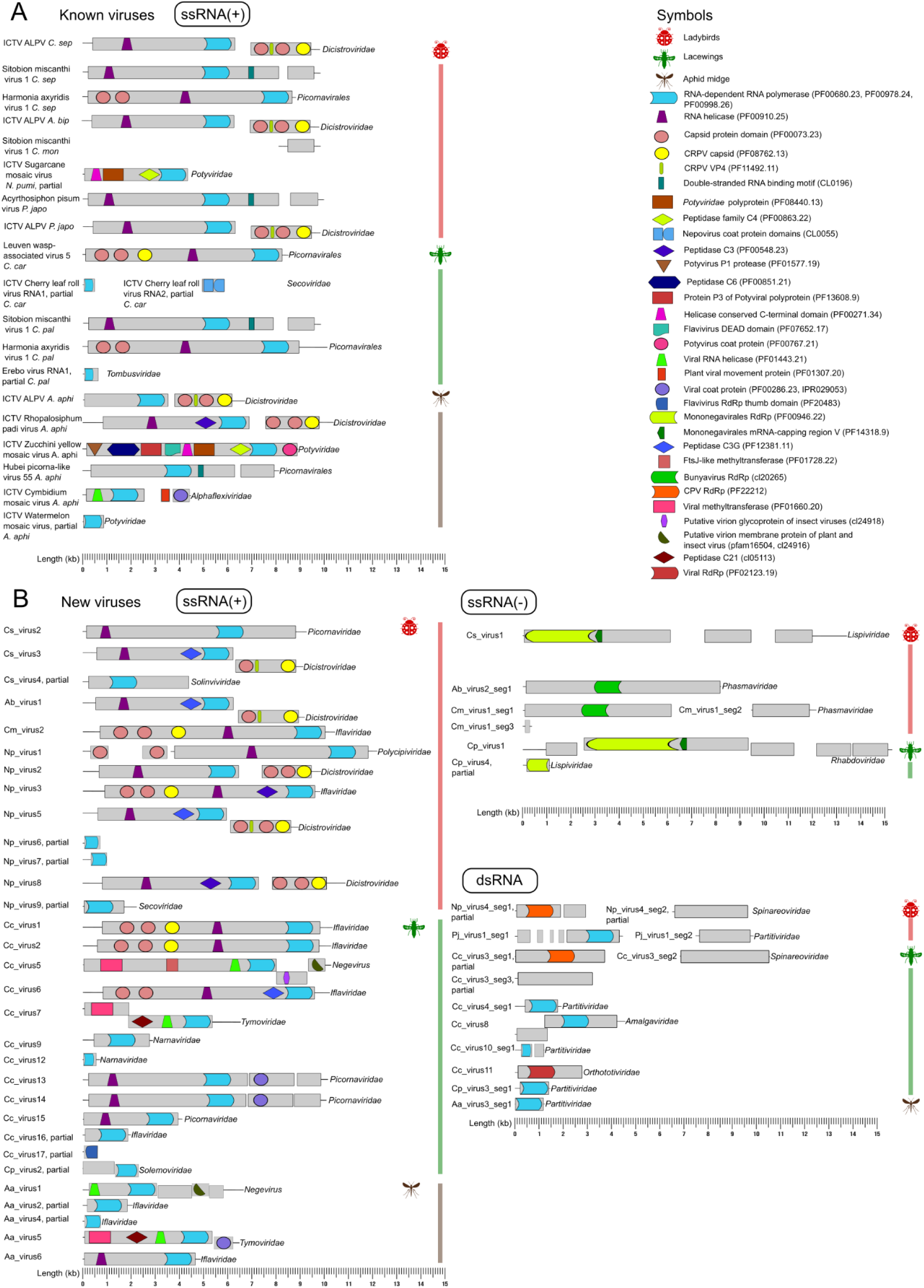
Representation of ORFs and domains of known and new viruses in scale (see ruler at the bottom), as well as information on their Baltimore classes and virus families. **A.** Known viruses found in ladybirds, lacewings, and aphid midges. **B.** Novel viruses detected in ladybirds, lacewings, and aphid midges. Red bars indicate viruses found in ladybird species, green bars indicate viruses of lacewing species, and light brown bars indicate viruses of the aphid midge.

**Table 2.** Overview of previously known viruses identified by sequence similarity searches.

| Contig/Location | Virus | Segment | Query size (nt) | Type | Query coverage (%) | Identity (%) | E-value | Closest sequence in GenBank | Original host/Country | GenBank accession |
| --- | --- | --- | --- | --- | --- | --- | --- | --- | --- | --- |
| Coccinella septempunctata (Coccinellidae) |  |  |  |  |  |  |  |  |  |  |
| known_virus_1 Germany | Aphid lethal paralysis virus isolate C. septempunctata | single | 9914 | nt | 94 | 96 | 0.0 | Aphid lethal paralysis virus isolate 4782ALPV | Apis mellifera, Spain | JX045858.1 |
| known_virus_2 China | Sitobion miscanthi virus 1 isolate C. septempunctata | single | 9942 | nt | 100 | 98 | 0.0 | Sitobion miscanthi virus 1 strain YYSMV1, complete genome | Sitobion miscanthi (aphid), China | MK733235.1 |
| known_virus_3 China | Harmonia axyridis virus 1 isolate C. septempunctata | single | 8833 | nt | 100 | 99 | 0.0 | Harmonia axyridis virus 1 | Harmonia axyridis, China | MK729664.1 |
| known_virus_4 China | Aphid lethal paralysis virus isolate C. septempunctata 2 | single | 6919 | nt | 79 | 98 | 0.0 | Aphid lethal paralysis virus isolate TIGMIC_1 | Haemaphysalis longicornis (tick), China | ON811696.1 |
| known_virus_5 China | Aphid lethal paralysis virus isolate C. septempunctata 3 | single | 5524 | nt | 100 | 99 | 0.0 | MAG: Dicistroviridae sp. isolate wag171dic2 | Motacilla cinerea (bird), China | MN918767.1 |
| Adalia bipunctata (Coccinellidae) |  |  |  |  |  |  |  |  |  |  |
| known_virus_6 Germany | Aphid lethal paralysis virus isolate A. bipunctata | single | 9915 | nt | 94 | 96 | 0.0 | Aphid lethal paralysis virus isolate 4782ALPV | Apis mellifera, Spain | JX045858.1 |
| Cryptolaemus montrouzieri (Coccinellidae) |  |  |  |  |  |  |  |  |  |  |
| known_virus_7 China | Sitobion miscanthi virus 1 isolate C. montrouzieri | single | 1342 | nt | 100 | 99 | 0.0 | Sitobion miscanthi virus 1 strain YYSMV1, complete genome | Sitobion miscanthi (aphid), China | MK733235.1 |
| Novius pumilus (Coccinellidae) |  |  |  |  |  |  |  |  |  |  |
| known_virus_8 China | Sugarcane mosaic virus isolate N. pumilus | single | 4397 | nt | 100 | 99 | 0.0 | Sugarcane mosaic virus isolate HZ7, complete genome | Zea mays (maize), China | JX047424.1 |
| Propylea japonica (Coccinellidae) |  |  |  |  |  |  |  |  |  |  |
| known_virus_9 China | Acyrtosiphon pisum virus isolate P. japonica | single | 10004 | nt | 99 | 99 | 0.0 | Acyrtosiphon pisum virus isolate APV-VV, complete genome | Acyrtosiphon pisum (aphid), China | MH301287.1 |
| known_virus_10 China | Aphid lethal paralysis virus isolate P. japonica | single | 9678 | nt | 99 | 95 | 0.0 | MAG: Aphid lethal paralysis virus isolate TIGMIC_1, complete genome | Haemaphysalis longicornis (tick), China | ON811696.1 |
| Chrysoperla carnea (Chrysopidae) |  |  |  |  |  |  |  |  |  |  |
| known_virus_11 Austria | Leuven wasp-associated virus 5 isolate C. carnea | single | 8529 | nt | 97 | 99 | 0.0 | Leuven wasp-associated virus 5 isolate Vvul_virus_34 hypothetical protein gene, complete cds | Vespula vulgaris (wasp), Belgium | MZ443602.1 |
| known_virus_12 Austria | Cherry leaf roll virus isolate C. carnea segment 1, partial | segment 1 | 556 | nt | 100 | 91 | 0.0 | Cherry leaf roll virus isolate Salzlandkreis-1_18 polyprotein 1 gene, complete cds | Weed, Germany | MN399680.1 |
|  | Cherry leaf roll virus isolate C. carnea segment 2, partial | segment 2 | 917 | nt | 100 | 91 | 0.0 | Cherry leaf roll virus isolate Salzlandkreis-1_18 polyprotein 2 gene, complete cds | Weed, Germany | MN399681.1 |
| Chrysopa pallens (Chrysopidae) |  |  |  |  |  |  |  |  |  |  |
| known_virus_13 China | Sitobion miscanthi virus 1 isolate C. pallens | single | 10255 | nt | 97 | 99 | 0.0 | Sitobion miscanthi virus 1 strain YYSMV1, complete genome | Sitobion miscanthi (aphid), China | MK733235.1 |
| known_virus_14 China | Harmonia axyridis virus 1 isolate C. pallens | single | 8903 | nt | 99 | 99 | 0.0 | Harmonia axyridis virus 1 | Harmonia axyridis, China | MK729664.1 |
| known_virus_15 China | Erebo virus isolate C. pallens segment 1, partial | segment 1 | 637 | nt | 99 | 94 | 0.0 | Erebo virus | Culex erythrothorax (mosquito), USA | MW434919.1 |
| Aphidoletes aphidimyza (Cecidomyiidae) |  |  |  |  |  |  |  |  |  |  |
| known_virus_16 USA | Aphid lethal paralysis virus isolate A. aphidimyza | single | 6741 | nt | 97 | 95 | 0.0 | Aphid lethal paralysis virus isolate 4782ALPV | Apis mellifera, Spain | JX045858.1 |
| known_virus_17 China | Rhopalosiphum padi virus isolate A. aphidimyza | single | 10358 | nt | 96 | 99 | 0.0 | MAG: Dicistroviridae sp. isolate ltt164dic1 | Bird, China | MN918744.1 |
| known_virus_18 China | Zucchini yellow mosaic virus isolate A. aphidimyza | single | 8904 | nt | 100 | 99 | 0.0 | Luffa aegyptiaca potyvirus strain pt075-pot-4 | Luffa aegyptiaca (plant), China | MN720007.1 |
| known_virus_19 China | Hubei picorna-like virus 55 isolate A. aphidimyza | single | 8149 | nt | 99 | 98 | 0.0 | Hubei picorna-like virus 55 strain spider134013 | Spiders, China | KX883691.1 |
| known_virus_20 China | Cymbidium mosaic virus isolate A. aphidimyza | single | 4457 | nt | 99 | 97 | 0.0 | Cymbidium mosaic virus | Plant, China | AM055640.2 |
| known_virus_21 China | Watermelon mosaic virus isolate A. aphidimyza, partial | single | 872 | nt | 100 | 98 | 0.0 | Watermelon mosaic virus isolate EM190182 | Squash, Spain | OQ680558.1 |

BlastN searches performed in October, 2024.

Overall, we identified 20 viruses (5 known, 15 new) in 5 ladybird species; 26 (5 known, 21 new) in 2 species of lacewings, and 11 (6 known, 5 new) in samples derived from the aphid midge. The lacewings showed the greatest abundance and diversity of viruses among the predators, with viruses classified into 13 families and 1 genus, while ladybirds carried viruses classified into 11 families. In samples derived from the unique aphid midge species assessed (*Aphidoletes aphidimyza*), viruses from 6 families and 1 genus (Figure 2 **A, B**) were detected. Worth noting, ladybirds shared 2 known viruses with lacewings (Sitobion miscanthi virus 1 and Harmonia axyridis virus 1); and 1 known virus with the aphid midge (Aphid lethal paralysis virus, ALPV) (**Table 2**, Figure 2A).

**Figure 2.**
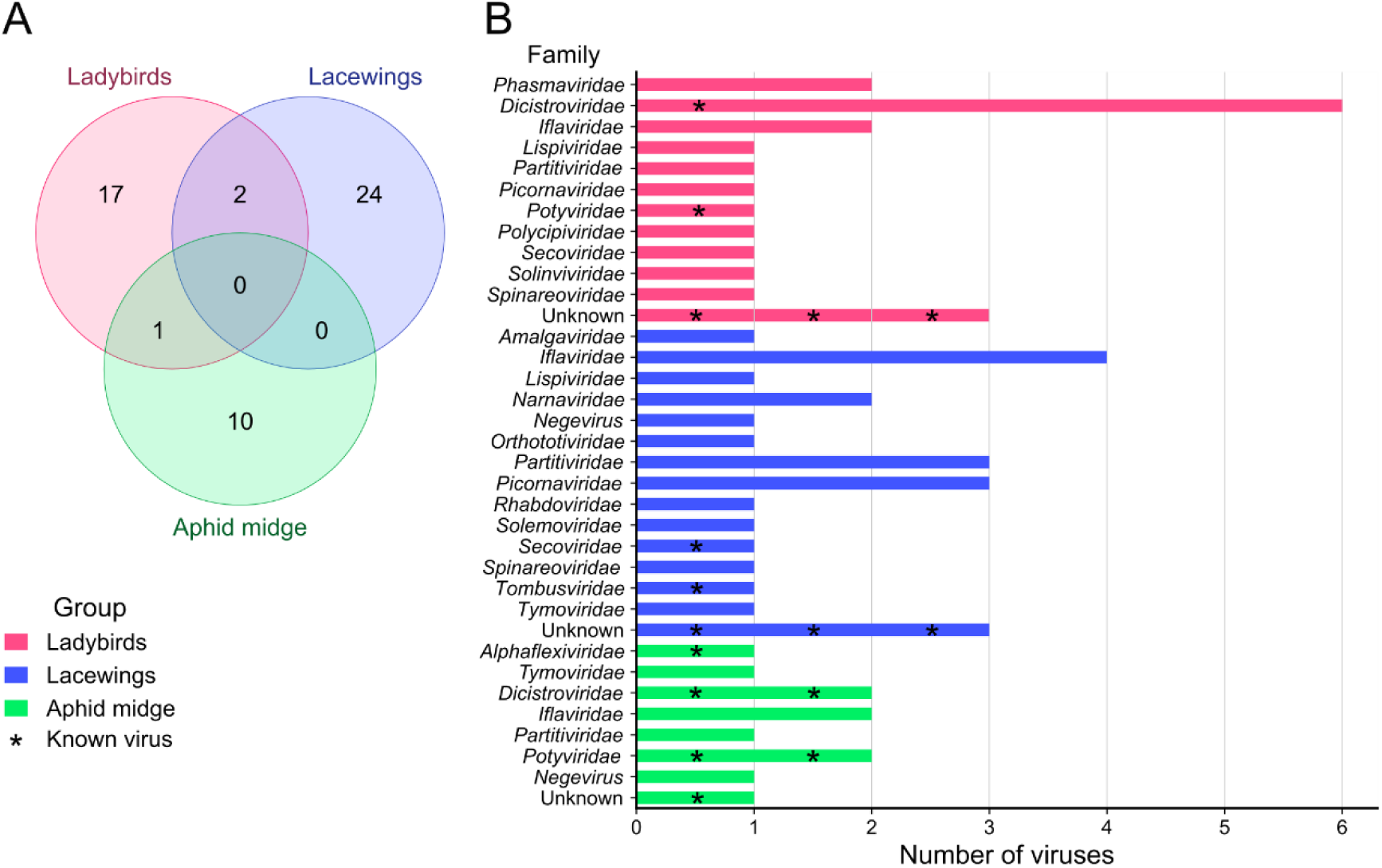
Distribution of viruses by each aphidophagous insect group and viral family. **A.** Venn diagram with the viruses found by each aphidophagous insect group. **B.** Bar graph showing the number of viruses found per viral family, and per insect group with corresponding colors. Known viruses are indicated with an asterisk over the bars.

### 3.2 Circulation of known viruses: potential new hosts

Viruses identified in aphids (and other hosts) have been identified in ladybirds (**Table 2**): ALPV and Sitobion miscanthi virus 1 were detected in *Coccinella septempunctata*; ALPV was detected in *Adalia bipunctata*; Sitobion miscanthi virus 1 was detected in *Cryptolaemus montrouzieri*, and ALPV and Acyrthosiphon pisum virus were detected in *Propylea japonica*. In addition, the plant pathogenic Sugarcane mosaic virus (SCMV, see **Table 2** for details) was detected in *Novius pumilus*. In lacewings, Leuven wasp-associated virus 5 and the plant pathogenic Cherry leaf roll virus were identified in *Chrysoperla carnea*. In addition, Sitobion miscanthi virus 1, Erebo virus, and Harmonia axyridis virus 1, a virus of the invasive ladybird *Harmonia axyridis* (also found in *C. septempunctata*, **Table 2**), were detected in *Chrysopa pallens*. For the aphid midge *A. aphidimyza,* aphid viruses (ALPV and Rhopalosiphum padi virus), plant pathogenic viruses (Zucchini yellow mosaic virus, Cymbidium mosaic virus, and Watermelon mosaic virus), and Hubei picorna-like virus 55 were also spotted (**Table 2**).

Additionally, the ALPV isolate C. septempunctata 3 here identified (**Table 2**), matched (at nucleotide level) with at least two *Dicistroviridae* sequences (GenBank: MN918767.1 and MN905993.1) with coverage >99% and identity >98% (e-value = 0.0). These ALPV sequences were isolated from cloacal swabs of two insectivorous wild bird species collected in China, *Motacilla cinerea* and *Poecile palustris* [82]. The researchers posited that *Dicistroviridae* and *Iflaviridae* viruses were widely present in wild birds compared to breeding birds, a reflection of the birds’ diet [82]. As a result, the highly similar strain of ALPV found in *C. septempunctata* and in those birds from China suggests virus detection in both trophic levels – ladybird (prey) and bird (predator). Another sequence (GenBank: MT138201.1) isolated from the cloacal swab of *Abrornis proregulus*, an insectivorous bird [83], presented 97% coverage and 98% identity at nucleotide level (e-value = 0.0) with the Sitobion miscanthi virus 1 isolate C. septempunctata and partial Sitobion miscanthi virus 1 isolate C. montrouzieri (**Table 2**). Finding the same virus in three trophic levels, to wit *Sitobion miscanthi* aphid (prey), *C. septempunctata* and *C. montrouzieri* (predators), and *Abrornis proregulus* (vertebrate predator) could suggest cross-species viral transmission (or at least dispersal) among trophic-related species. For an overview, Figure 3 illustrates a Sankey diagram with the 13 known viruses found in each invertebrate predator species and in other organisms (which were determined by sequence similarity search best hits at nucleotide level, see **Supplementary Table 3**).

**Figure 3.**
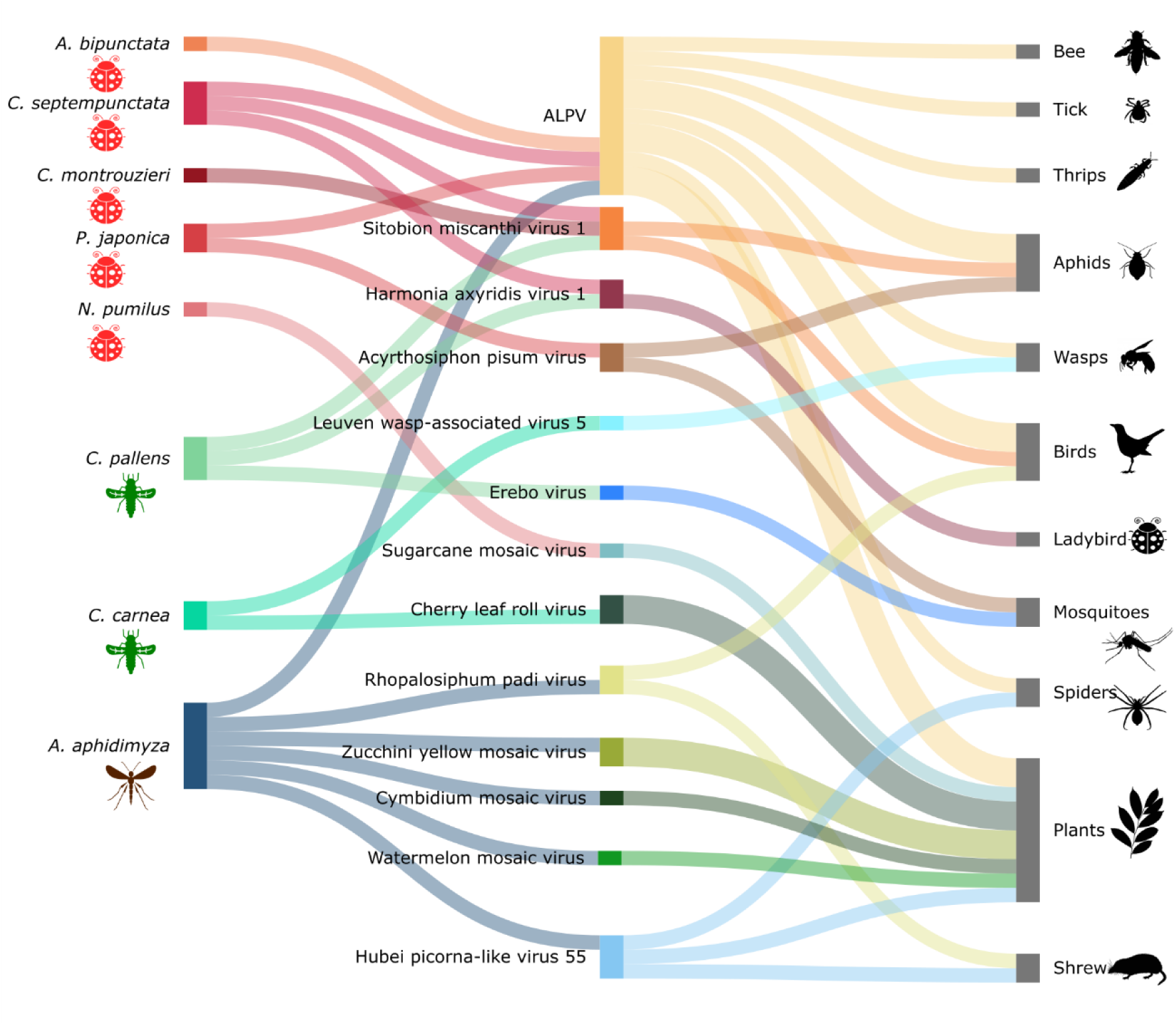
Sankey plot showing the known viruses and their respective associated eukaryotic species. On the left, the predatory species here investigated, to wit 5 ladybird species (red symbol), 2 lacewing species (green symbol), and 1 aphid midge species (brown symbol). On the right, other organisms in which the known viruses were detected, revealed by BlastN searches on NCBI (black symbols). ALPV: Aphid lethal paralysis virus.

### 3.3 Phylogenetic inference of new viral sequences

Transcriptomic assemblies yielded 41 putative novel viral sequences (see **Table 3** for details) encoding the RdRp domain (Figure 1B). Of these, 27 genomes are complete or nearly complete compared to the size of the closest viral sequences. The other 14 sequences represent partial genomes, including viruses that are segmented, such as reoviruses. The 41 sequences were subjected to phylogenetic analysis, and the virus nomenclature was determined by its taxonomic classification and the name and/or characteristics of the host in which it was found. All phylogenetic trees are available in the supplementary materials. The taxonomic classification and Latin binomial names of the new viruses are shown in Figure 4.

**Table 3.**
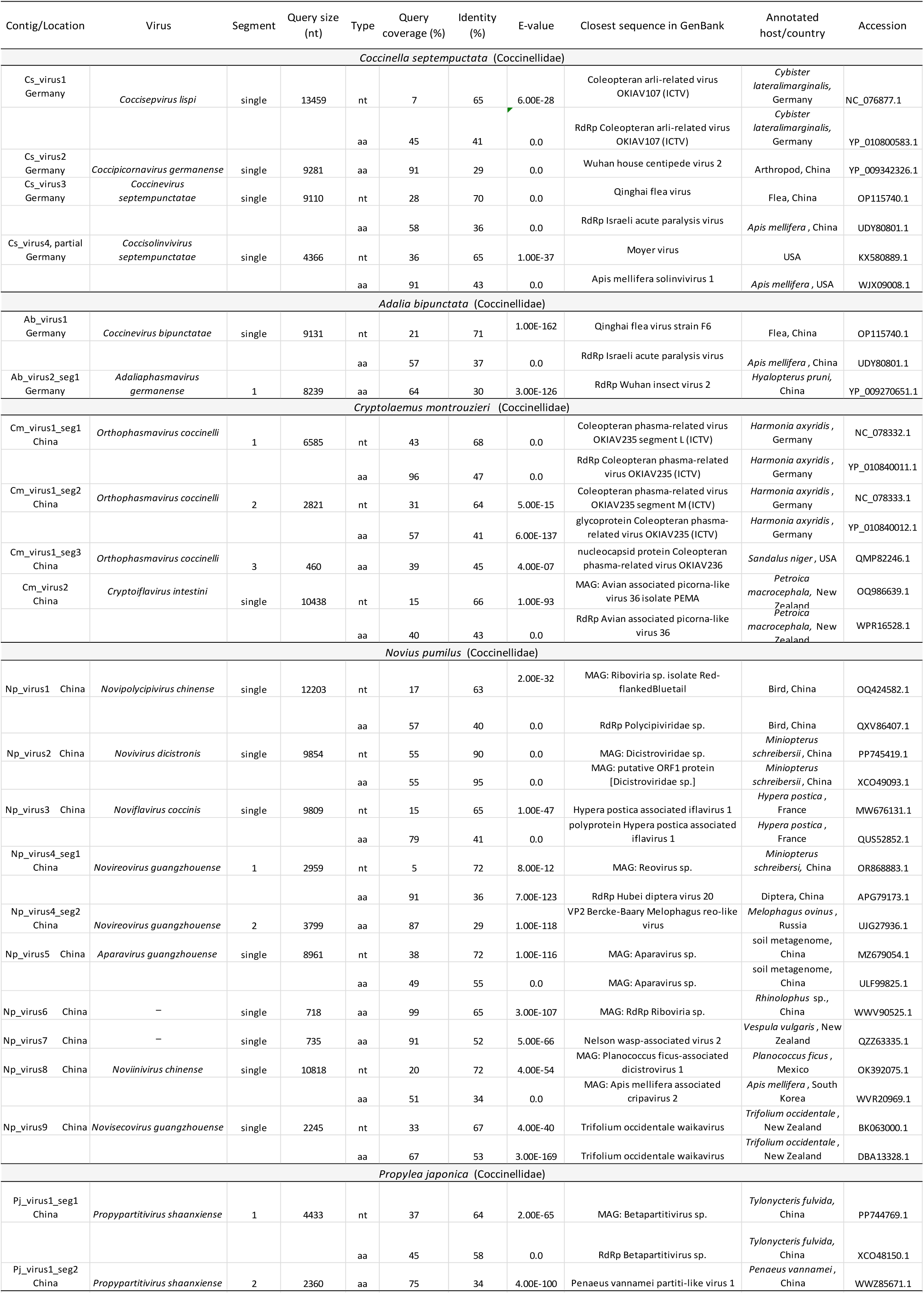
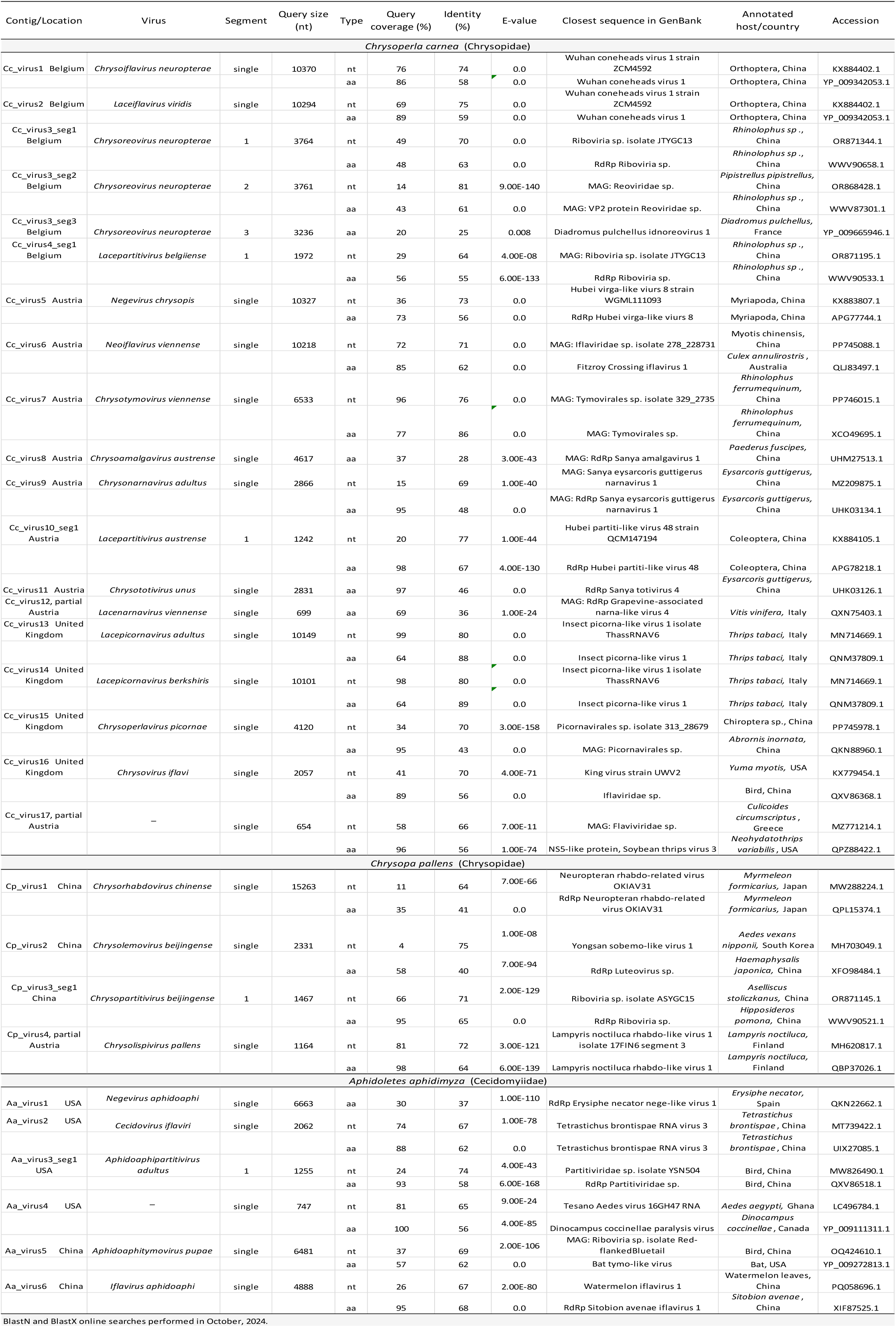
Overview of new viruses identified by sequence similarity searches.

**Figure 4.**
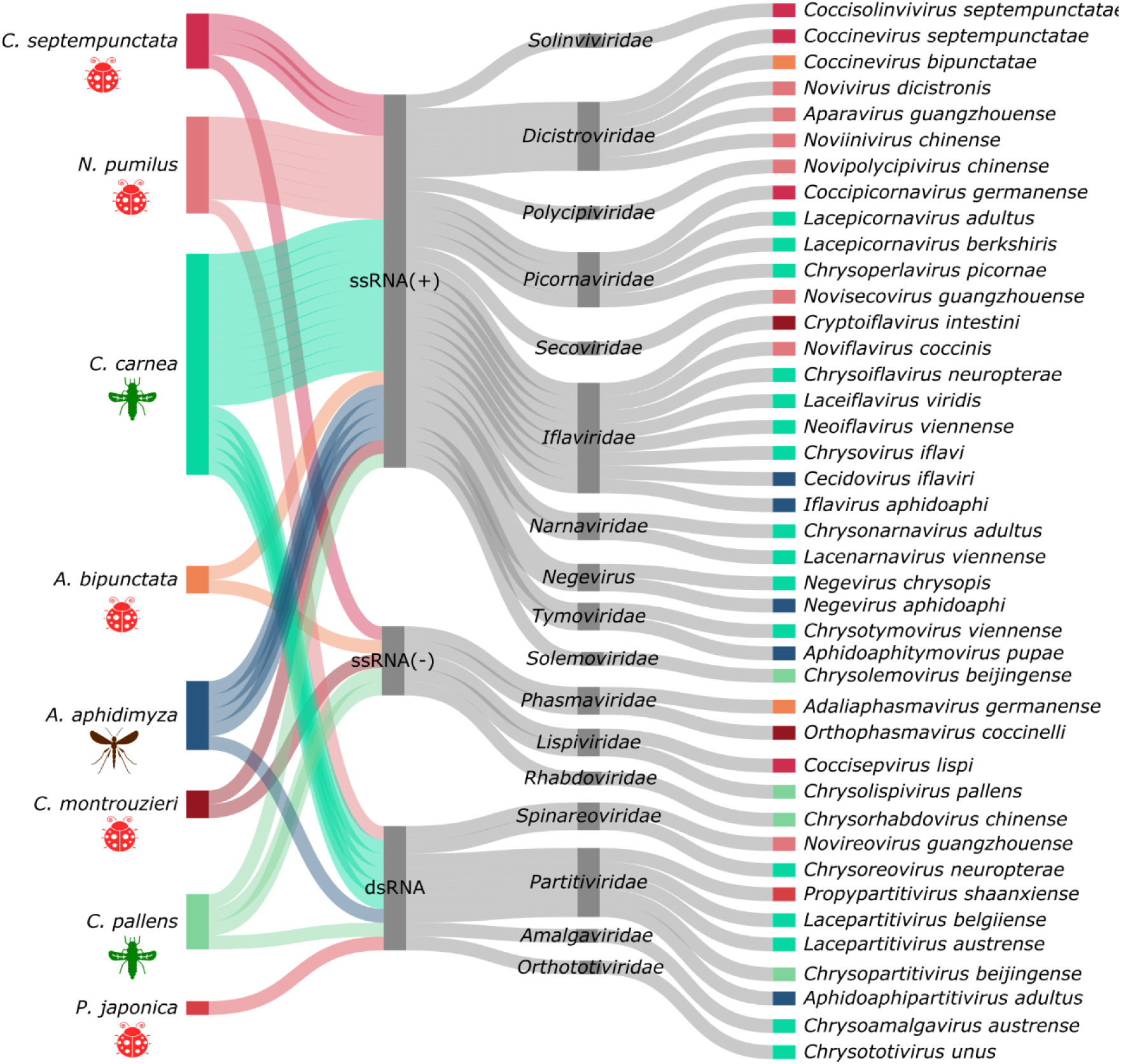
Sankey plot showing the Baltimore and taxonomic classifications followed by the Latin binomial names of the new viruses for each predatory species. On the left, the description of the predatory species investigated; ladybird species are represented in reddish colors, lacewings in greenish colors, and the aphid midge is represented in blue; in the middle, the three Baltimore classes into which the viruses were classified, and the description of 16 viral families and 1 genus obtained from the phylogenetic analyses; on the right, the proposed binomial names for each virus with rectangles corresponding to its associated predator color.

### 3.4 Relatedness of new viruses with known viral species

The sequence similarity searches revealed the closest sequence in GenBank for each new virus detected (**Table 3**). The examination of phylogenies and respective hosts may offer insights into the evolutionary and ecological trajectories of the viruses in question. The closest viruses identified by blast searches have been found in prey (such as aphids, mealybugs, and thrips), invertebrate predators (such as wasps and myriapods), vertebrate predators (several insectivorous bats and birds), ectoparasites (such as fleas, flies and ticks), as well as other insects (such as beetles, honeybee, mosquitoes, and orthopterans). In addition, a few new viruses closely resembling plant viruses were registered (see **Table 3** for details). A notable finding was the presence of viruses, in aphid-consuming insects, that are closely related to viruses of organisms at higher trophic levels, such as bats and birds. The *Novivirus dicistronis,* isolated from *N. pumilus,* grouped with a dicistrovirus (XCO49093.1) hosted by a bat (*Miniopterus schreibersii,* 100% bootstrap confidence, Figure 5A). Furthermore, iflaviruses detected in ladybirds and *C. carnea* are positioned in close proximity to viruses belonging to vertebrate predators, in addition to the insect viruses positioned in the tree (Figure 5B). Two picornaviruses of *C. carnea* (*Lacepicornavirus adultus* and *Lacepicornavirus berkshiris*) are in the same clade (100% bootstrap) of a picornavirus (QNM37809.1) isolated from its prey, *Thrips tabaci* (Figure 5C). A tymovirus isolated from *C. carnea* (*Chrysotymovirus viennense*) clustered with a virus (XCO49695.1) from a bat (*Rhinolophus ferrumequinum*, 100% bootstrap). Also, *Aphidoaphitymovirus pupae,* found in *A. aphidimyza*, clustered with another virus from a bat (YP_009272813.1, 85% bootstrap), and it is in the same clade as a bird virus (WKV33971.1, 100% bootstrap, Figure 5D). Partitiviruses found in the predators evaluated clustered into more diverse clades, which included viruses from birds, bats, shrew, pigs, flies, dragonflies, and more (Figure 5E). *Lacepartitivirus belgiiense* grouped with a virus from a bat (WWV90533.1, *Rhinolophus* sp.) with 100% bootstrap support (Figure 5E). In the *Spinareoviridae* tree, there are viruses from mammals as well, with *Chrysoreovirus neuropterae* grouped with a virus isolated in a bat (WWV90658.1, *Rhinolophus* sp.) with 100% bootstrap (Figure 5F). In addition (see supplementary materials), *Novipolycipivirus chinense*, a virus isolated from *N. pumilus*, clustered with a bird virus (QXV86407.1, 100% bootstrap). However, the rhabdovirus *Chrysorhabdovirus chinense*, found in *C. pallens*, was grouped with a neuropteran virus (100% bootstrap).

**Figure 5.**
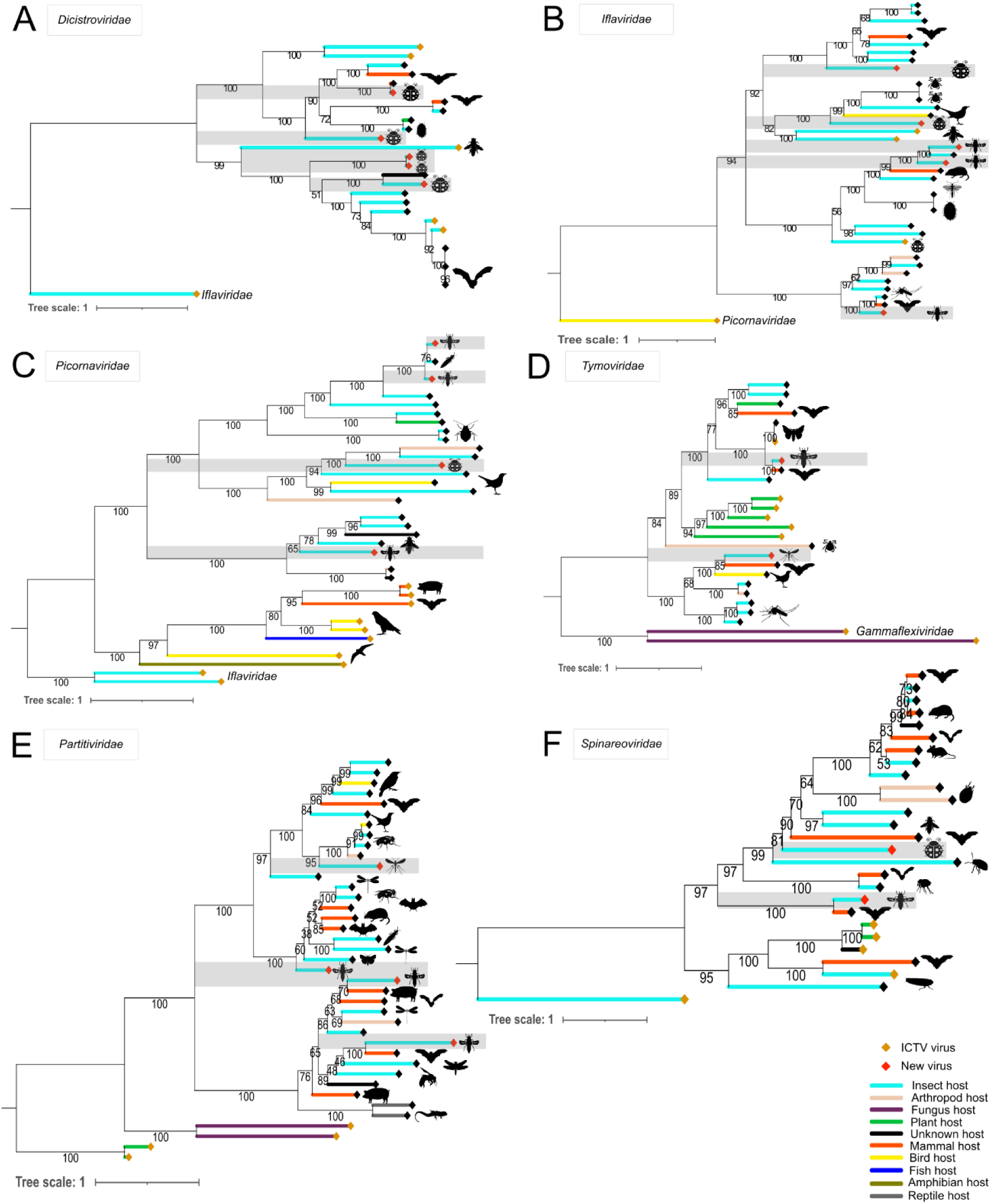
Phylogenetic trees showing new viruses and their respective species of origin, as well as the species of origin of closely related virus species. **A.** *Dicistroviridae* viruses; **B.** *Iflaviridae* viruses; **C.** *Picornaviridae* viruses; **D.** *Tymoviridae* viruses; **E.** *Partitiviridae* viruses, and **F.** *Spinareoviridae* viruses. The novel viruses described in this study are highlighted in grey. The lines of the trees are colored according to the higher taxonomic classifications of the possible host of each virus (see the legend at the bottom right of the figure).

### 3.5 Abundance and geographical distribution of the viruses

The expression levels of constitutive genes (*calmodulin* and *histone H3*) and the viral abundance, in transcripts per million (TPM), were determined for all the libraries included in the study (see **Supplementary Table 4**). Figure 6 presents a heatmap with the normalized values of quantification log10 (TPM+1), as well as the representation of libraries divided by geographical origin. Both species of lacewings displayed divergent viral profiles, exhibiting variation in their viral composition according to the geographical location. Yet, *Lacepartitivirus belgiiense* (Cc_virus4_seg1) was quantified in three libraries of *C. carnea* from different countries (Austria, Belgium, and UK). Sitobion miscanthi virus 1 and Harmonia axyridis virus 1 were found in high levels in *C. pallens* from China (Figure 6). The five species of ladybirds had different patterns of virome composition, and libraries of *C. septempunctata* from China and Germany contained only ALPV in common, in high levels (Figure 6). ALPV was also found in high levels in *P. japonica* (China) and *A. bipunctata* (Germany). Finally, two libraries of *A. aphidimyza* from different countries also exhibited different viruses. For instance, *A. aphidimyza* from China had Rhopalosiphum padi virus in high abundance, while the same species from USA exhibited ALPV at a high level (Figure 6). Consequently, ALPV was identified as the most prevalent and abundant virus in this analysis, given its presence in four species of predators and three countries.

**Figure 6.**
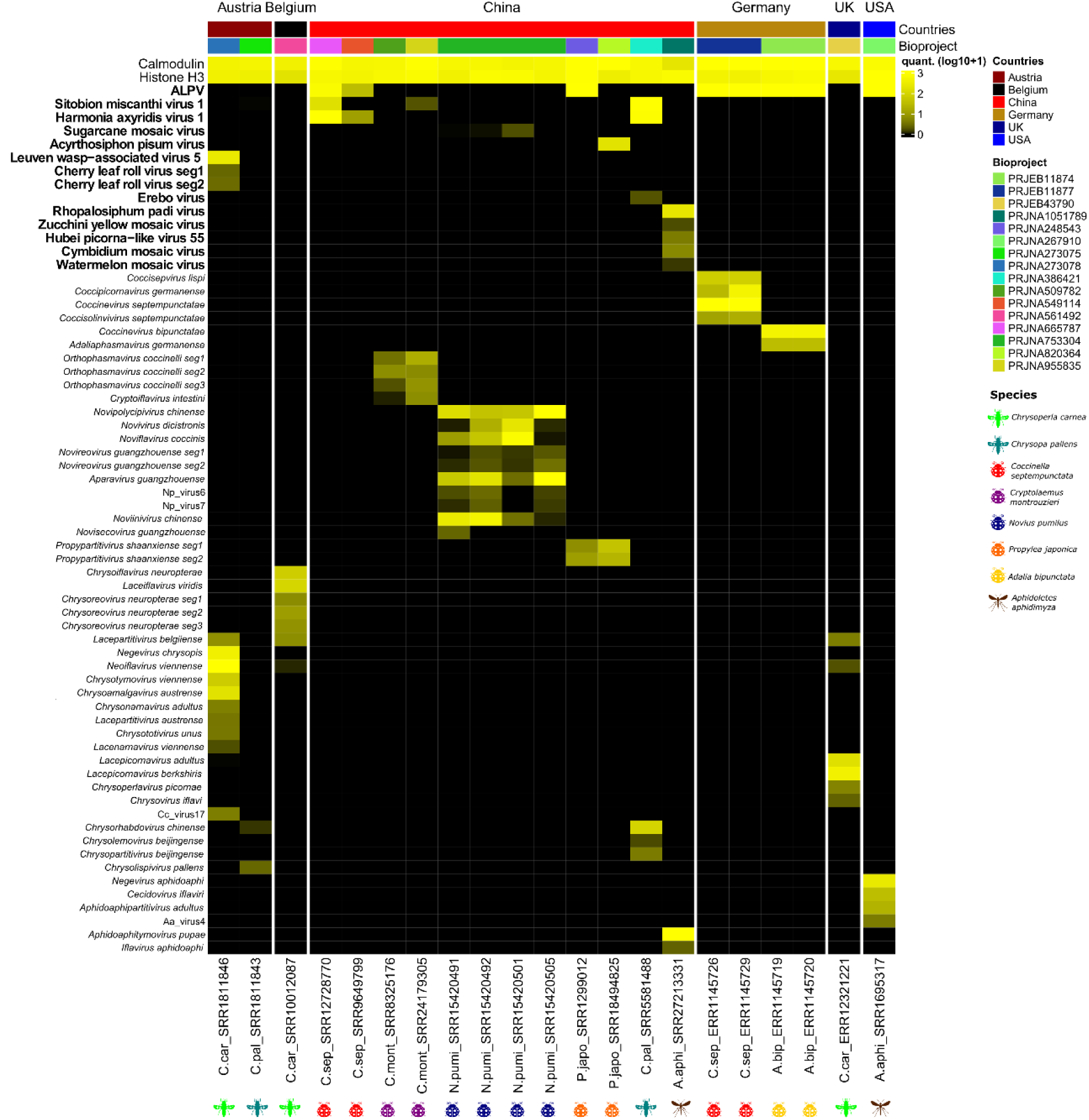
Heatmap showing the transcript abundance of the viral sequences and constitutive genes estimated in transcripts per million (TPM). The known viruses are highlighted in bold. Above, six countries and their bioprojects are represented in different colors. Underneath, the RNA-seq libraries utilized are represented with colored symbols by predator species.

## 4. Discussion

In this study, it was investigated which viruses are present in 21 publicly available RNA deep sequencing libraries from 8 predatory insect species that control agricultural pests globally. As far as we know, this is the first study focused on the investigation of viruses in three groups of aphidophagous insects used in the biological control of soft-bodied insects. In addition to the need to identify and monitor pathogens that could threaten the survival of important biological control agents, it is known that microbial symbionts may influence the ecology and evolution of host species, and consequently on the dynamics of trophically-related organisms, as predator and prey [84,85]. Therefore, research on viruses that circulate in agricultural insect species is fundamental for improving biological control strategies based on trophic interactions, for biodiversity conservation, and for developing knowledge on the evolution of viral entities and their role as members of an agroecological system.

### 4.1 Are the viruses from the prey actually infecting the predators?

Research on viruses infecting predatory ladybirds, lacewings, and aphid midges is not prevalent. Therefore, the corpus of knowledge on this subject is in its infancy and many biological questions will need to be answered with further studies. However, it is imperative to consider the third-party sample preparation process for RNA transcriptomes, and the biological context of the species, when available, in order to gain insights into the viruses detected in this analysis. Details on the libraries and predatory species are available in **Table 1**. The libraries were selected randomly by species, yet, most of the samples used for RNA extraction were from laboratories. Given the well-established rearing conditions (mentioned below), the origin of the viruses in question can be partially understood. In addition, the viral abundance may help to understand the prevalence of the virus in its associated host (Figure 6). However, the absolute absence of virome studies involving the species analyzed, in different populations and conditions, makes it very difficult to establish the clear origin or replicative capacity of the viruses found. In addition, the need for several experimental studies to understand the infection or replication of a virus in a given host is well understood, but such studies are focused on viruses of medical interest. Therefore, this question is likely to remain open for some time.

Viruses identified in specimens that were left under starvation for some hours, before further processing, are more likely to belong to them instead of their prey, since this caution is an assumption that remains of the ingested prey have been eliminated, reducing the likelihood of viral contamination. In addition, fed specimens may actually be infected with a “prey virus” and this cannot be clarified in this study. The fourth-instar larvae of *C. septempunctata* used in SRR9649799 were left for 12h under starvation before RNA extraction [86]. *C. septempunctata* adults from SRR12728770 were maintained in laboratory supplied with *Aphis glycines* daily [87]. *C. septempunctata* adult samples of PRJEB11877 (Figure 6) were collected in Germany and maintained in laboratory with bean plants infected with pea aphids (*Acyrthosiphon pisum*) [88]. Adult samples of *A. bipunctata* were obtained from a commercial source and maintained under identical conditions [88]. Fourth-instar larvae of *C. montrouzieri* (SRR8325176) were left for 12h under starvation before RNA extraction [47]. The four RNA-seq libraries of *N. pumilus* were obtained from the same bioproject (Figure 6). Adults of N. pumilus were fed with *Icerya aegyptiaca* (Hemiptera, Monophlebidae) or not fed (i.e. left under starvation) for 24h [89]. Here, 4 libraries of *N. pumilus* (3 fed, 1 under starvation) were analyzed, and the viruses detected were the same, only with variation in abundance (Figure 6), except for Np_virus6 and Np_virus7, that did not appear in the library of the sample under starvation (SRR15420501). Interestingly, these contigs were considered as “fragments of viral origin” in this analysis due to their sizes (<1000nt, **Table 3**). Consequently, they are probably digested virus fragments obtained from food, which can explain why the ladybird under starvation does not have them. Another virus that appeared once in a library of fed *N. pumilus* was the Np_virus9 (*Novisecovirus guangzhouense*, 2245nt, Figure 6). In this case, further research is needed to determine whether the virus is from the prey (i.e., the food source) or the predator, or both. Another interesting finding is the abundance of Sugarcane mosaic virus in libraries of fed ladybird (0, 0.010 and 0.034 TPM) *versus* under starvation (0.727 TPM, **Supplementary Table 4**). A hypothesis for the Sugarcane mosaic virus’ higher abundance (21 times more abundant) in the ladybird under starvation (SRR15420501, Figure 6) is that it is more vulnerable to the infection because it is under stress, thus the virus could replicate more easily, however, the possibility of Sugarcane mosaic virus infecting ladybirds needs to be assessed. The laboratory samples of *P. japonica* (SRR1299012) were maintained on *Aphis craccivora* for more than two years [90]. Pj_virus1_seg1 and Pj_virus1_seg2 (*Propypartitivirus shaanxiense*) appeared in two libraries of *P. japonica* from distinct bioprojects (Figure 6), but it is not possible to ensure this virus belongs to *P. japonica* or its prey. Specimens of *C. carnea* and *C. pallens* from Austria were collected, preserved in RNAlater, and stored at 4°C until RNA extraction and further processing [91]. In addition, specimens of *C. carnea* were purchased from a company in Belgium, and maintained with *Myzus persicae* aphids until further processing [92]. Details on the sample of *A. aphidimyza* are discussed in the next session.

### 4.2 The known viruses and their potential ecological repercussions

Of thirteen known viruses here detected in aphidophagous insects, 7 (54%) have already been found in aphids. ALPV was found to be expressed at levels 5.8 times higher than *calmodulin* in *C. septempunctata*, 1.3 times higher in *P. japonica*, and 2.6 times higher in *A. aphidimyza*. *Calmodulin* expression levels in *A. bipunctata* were found to be 1.38 times higher than those of ALPV, yet the virus was detected at a concentration 2.1 times higher than the constitutive *Histone H3* in this species (**Supplementary Table 4**). ALPV is a *Cripavirus* of *Dicistroviridae* that infects different aphid species, which exhibit variable pathological symptoms in response to it (reviewed in [18]). Because ALPV can cause paralysis and uncoordinated movements [93], it is suggested that ALPV-positive (ALPV+) aphids are more vulnerable to predation, which would justify the high prevalence and abundance of ALPV in the predators evaluated in this study (8/21 RNA-seq libraries were ALPV+). This hypothesis is consistent with the experimental evidence obtained with aphids infected by another dicistrovirus, Rhopalosiphum padi virus (RhPV), that causes pathological interferences as well, in a study conducted with infected and uninfected *Rhopalosiphum padi* aphids [94]. Ban *et al*. (2008) tested the attack rate of predation by two biological control agents, *Coccinella septempunctata* (ladybird) and *Aphidius ervi* (parasitoid wasp), and found that the mean of infected aphids (RhPV+) consumed by *C. septempunctata* adults, in 24 hours, was higher than that of uninfected aphids [94]. Moreover, parasitoids attacked more RhPV+ aphids than uninfected ones. The authors argued that RhPV+ aphids had weaken behavioral defenses, consequently, they became susceptible prey to capture [94]. In this study, RhPV was found only in 1 library of *A. aphidimyza* from China, and this finding may be related to the aphid species made available for predation in the laboratory breading. RhPV also has been found in vertebrate predators, such as birds and shrew (Figure 3), suggesting its intertrophic dispersal or transmission. Worth noting, in addition to ladybirds and the aphid midge, ALPV and its variants have been found in several species (whether prey or predator, Figure 3), to wit diverse aphid species of economic importance (reviewed in [18]), whitefly [95], bees and wasps [96–98], beetles [99], spider and damselfly [100], bats [101], birds [82], and plants [102,103]. Therefore, the apparent ease with which ALPV can access or infect possible host species at different trophic levels is a strong indication that the virus may be transmitted by ongoing exposure through the feeding, and trophic relationships, of the species described as hosts to date. Thus, a rigorous investigation is warranted to ascertain whether the infection of prey (hemipterans, for example) by ALPV confers a competitive advantage to specific predators, such as ladybirds, by facilitating hunting. This inquiry is of particular relevance, as a natural virus that does not cause disease in plants, but weakens the pest could be employed to enhance the efficacy of biological control by predatory insects. However, it is imperative that the predator’s exposure to the same virus is asymptomatic or does not result in any adverse impact on its biological performance. The development of effective and safe methods for the combined use of naturally occurring organisms (such as predator + virus) for the biological control of agricultural pests is of vital importance. This necessity arises due to numerous ecological problems caused by the overuse of pesticides [104], and climate change, that can impact on the efficiency of biocontrol agents, since the global warming may increase the distribution and abundance of pests [105].

There are harmful, commensal (or neutral), and mutualistic (those that can increase host fitness) viruses infecting insects [106], and these infection strategies are changeable according to a multitude of factors [107]. Acyrthosiphon pisum virus (APV) is a picorna-like virus that infects the aphid *Acyrthosiphon pisum*, thereby reducing the development and growth of its host [108]. It is transmitted to plants during aphid feeding, but it is not phytopathogenic. However, mutations in APV RdRp increased the replication of the virus in the aphid, which helped suppress the plant’s defenses against aphids. Thus, the virus enabled the aphid to adapt to new plants that were previously incompatible with the aphid’s development [109]. The suppression of the plant’s defenses by the APV is due to a reduction in jasmonic acid phytohormone levels, which favors the aphid’s survival [110]. APV was found in 1 library of *P. japonica* ladybird, with 200 TPM concentration, but its effects on the predator are unknown. Sitobion miscanthi virus 1, another aphid virus, has been found in libraries of two ladybirds and one lacewing (Figure 6). In *Chrysopa pallens*, a lacewing that exhibits predatory behavior throughout its entire lifespan (Figure 7A), this virus is found up to 35 times more expressed than constitutive genes (**Supplementary Table 4**), nonetheless, its effects on predators and aphids remain to be elucidated. As an example of the relevance of this topic, a study demonstrated that lacewings *Chrysoperla externa* fed with aphids infected with the potato leafroll virus (*Solemoviridae: Polerovirus*) had a prolonged larval period and reduced survival rate compared to lacewings fed with uninfected aphids [111].

**Figure 7.**
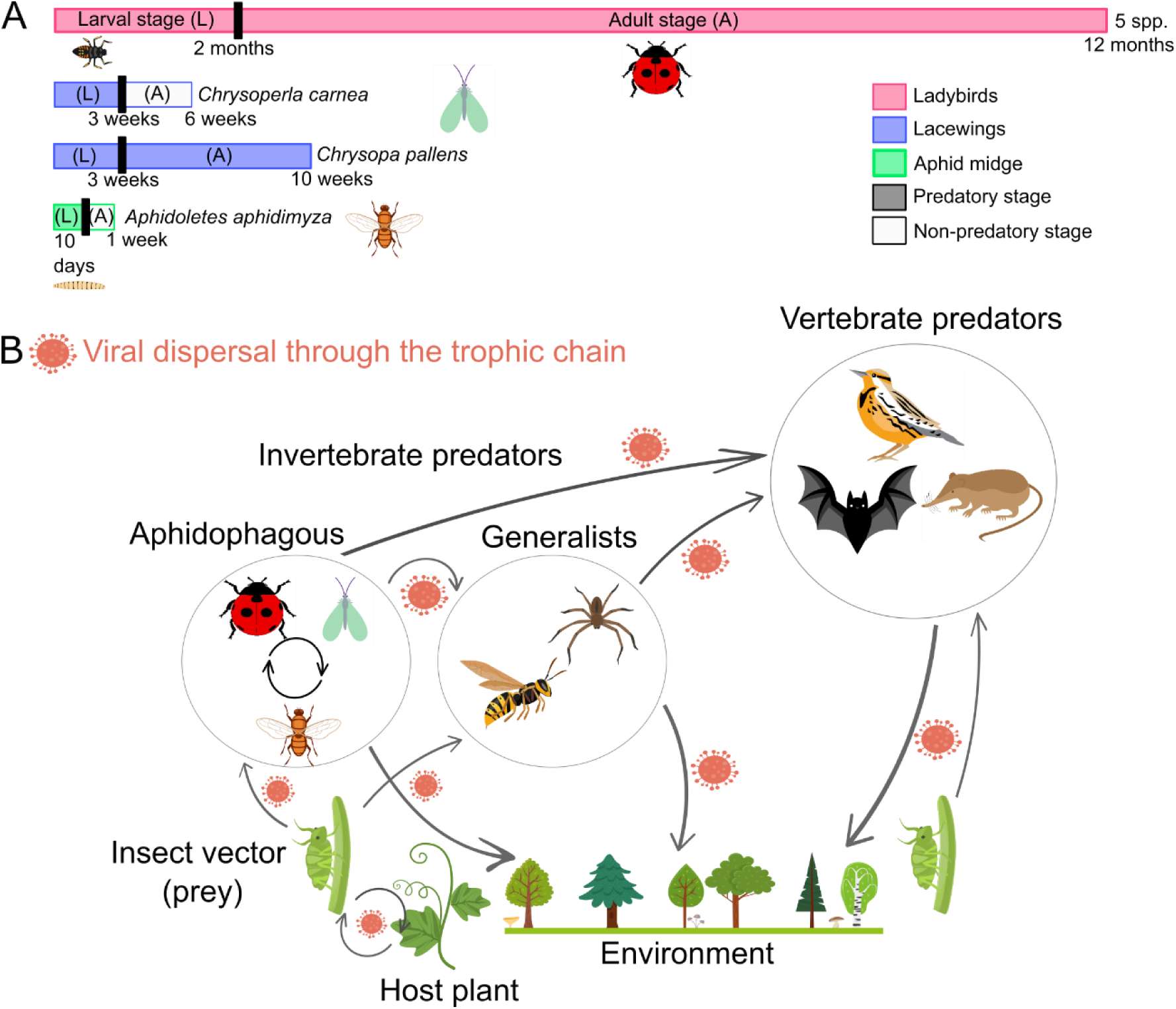
**A.** Representation of the predatory and non-predatory stages of the aphidophagous insects included in this study. Ladybirds and *C. pallens* are predatory in both the larval and adult stages; *C. carnea* and *A. aphidimiza* are only predatory in the larval stage. The pupal stage was not represented because it is a non-feeding stage. **B.** Simplified model of possible routes of viral dispersal/transmission among trophic-related species. There are representatives of insect prey (or pests), invertebrate predators (aphidophagous or generalists), and vertebrate predators (such as birds, bats, and small mammals).

Furthermore, three phytopathogenic viruses have been found in predators and are vectored by aphids. Sugarcane mosaic virus, Zucchini yellow mosaic virus (ZYMV), and Watermelon mosaic virus (WMV) have been found in lower abundances than the expression of constitutive genes (Figure 6, **Supplementary Table 4**). Finding phytopathogenic viruses in aphidophagous predators suggests that, in addition to the successful elimination of the pest insect, a cycle of transmission of the pathogenic viruses by the vector to new plants has been disrupted. Plants are sessile and have a cell wall as a protective barrier, consequently, viruses that infect plants rarely are transmitted through direct contact in natural environments [112]. Instead, plant viruses predominantly rely on vectors to be transmitted [112]. Because ladybirds and pupae of aphid midges do not act as viral vectors (such as hemipteran insects, that use their mouthparts like needles to drill and suck sap or the plant cell’s content), those pathogenic viruses are not going to infect new plants. Given the presence of these viruses in low concentrations in the predators, it is plausible that they do not exert deleterious effects. However, further research is necessary to ascertain the infection and extent of any potential adverse consequence. In plants, SCMV [113], ZYMV [114], and WMV[115] are devastating pathogens.

The other six known viruses found in the aphidophagous predators have not been found in aphids. Cymbidium mosaic virus (CMV) and Cherry leaf roll virus (CLRV) are pathogenic viruses of orchids [116] and cherry (as well as other fruit plants) [117], respectively. A biological vector for these viruses is unknown [116,118]. Since the *C. carnea* library that contains CLRV was sequenced with adult specimens, which are not predatory [6], it is suggested that the virus was caught from infected plants (through nectar or pollen ingestion). However, the *A. aphidimyza* pupae used for construction of the library that contains CMV (Figure 6, **Table 1**), and other viruses (RhPV, ZYMV, Hubei picorna-like virus 55, and WMV) was reared in laboratory, preying on aphids (*Aphis craccivora*, hosted by broad bean seedlings in insect-rearing cages of a glass greenhouse) [119]. Because the pupal stage is a non-feeding period [120] (in *A. aphidimyza* it lasts 8-12 days [119]), those viruses were caught by the insect in its larval predatory stage (Figure 7A) or by vertical transmission. Thus, if the source of infection was the aphid, the potential of *Aphis craccivora* as a vector for CMV (and/or as vector to the other viruses identified in pupae of *A. aphidimyza*) should be investigated. Erebo virus (EV) and Leuven wasp-associated virus 5 (LWaV5) were identified for the first time in 2021 by transcriptomic studies [121,122]. EV was isolated from mosquitoes (samples of *Culex* spp. and *Culiseta* sp.) collected in California, USA [121], and LWaV5 was identified from *Vespula vulgaris* larvae from Belgium [122]. In this research, EV and LWaV5 were detected in two lacewing species, *C. pallens* (China), and *C. carnea* (Austria), respectively (Figure 6). As both viruses were detected in predators from countries other than those where they were first identified (USA and Belgium), it is possible to infer that they are circulating in a wider geographical area than the regions initially investigated. Moreover, Hubei picorna-like virus 55 was identified in 2016 from spiders collected in China [123]. Since then, it has been identified in other species such as prey (soybean thrips [124]), and predators (other spider species [100], shrew [125], and the aphid midge here investigated, all from China).

The harlequin ladybird (*Harmonia axyridis*) is native to Asia; however, it is regarded as an invasive species on a global scale [126]. This species poses a threat to biodiversity, especially among aphidophagous insects. Several countries have reported detrimental effects on native species, primarily coccinellids, due to competition and predation [126]. *H. axyridis* hosts pathogens (such as bacteria, fungi, and protists), and parasites [127,128], but there are few viruses identified from it in the literature. Here, Harmonia axyridis virus 1, a virus identified in 2020 from *H. axyridis* adults collected in China [54], was detected in 2 libraries of *C. septempunctata* from China, and 1 library of *Chrysopa pallens* (lacewing) also from China. In both species, the virus was found in concentrations higher than those of the constitutive genes (Figure 6, **Supplementary Table 4**). Thus, in addition to the ecological problems caused by the invasive species, there is a risk of transmitting microorganisms (e.g. viruses) that can have unknown effects on native aphidophagous insects. *Harmonia axyridis* is a well-studied example that demonstrates invasive species boast superior immune systems in comparison with evolutionarily related species that are not invasive [129]. This is explained by exposure to pathogens and parasites encountered in newly inhabited environments [129]. Briefly, *H. axyridis* pre-injected with bacteria and yeast cells yielded an immune response with the expression of around 50 genes encoding inducible antimicrobial peptides (AMPs), the maximum number documented at that time for any animal species [130]. Also, *H. axyridis* is more resistant to pathogens that kill native ladybirds, such as *A. bipunctata* and *C. septempunctata* [129,130]. In light of these circumstances, the absence of surveillance for invasive ladybird viruses, such as those of *H. axyridis*, in countries facing threats from invasive species, constitutes a significant concern. Furthermore, the lack of knowledge on viruses present in native species also serves as a hindrance to the effective utilization of aphidophagous insects as biological controllers, given that ladybirds and lacewings can share viruses in common (such as Harmonia axyridis virus 1). Worth noting, the effect of Harmonia axyridis virus 1 infection on its hosts is unknown.

### 4.3 On the diversity of new virus discoveries

The 41 new viruses were classified into 16 families and 1 genus (Figure 2B). Further details concerning the viral families are provided in the supplementary text. It is therefore remarkable that aphidophagous insects can harbor a great diversity of viruses. In this study, the utilization of 21 RNA-seq libraries from only eight species of predatory organisms resulted in the identification of 41 putative new viruses and 13 known viruses. However, given the number of predatory species, this is merely a minuscule fraction of the total number of viruses associated with these organisms and trophic-related species that compose an agricultural ecosystem. There are around 6000 recorded species of Coccinellidae, more than a hundred with great potential as biocontrol agents [131]. Approximately 1200 species of Chrysopidae are extant globally [132]. On Cecidomyiidae, there are currently 6651 known species, although the number of species of this family is immeasurable [133]. Consequently, to further advance understanding of viruses, especially in terms of their biodiversity, and to optimize biocontrol mechanisms involving viruses in ecological systems, additional research is necessary.

### 4.4 Dispersal/transmission routes and the presence of common viruses in different trophic levels – many uninvestigated possibilities

To summarize the results obtained in this analysis of viruses in aphidophagous insects, a network of ecological interactions among organisms of different trophic levels was created. Along with trophic interactions, possible routes of virus dispersal are represented (Figure 7B). It is clear that trophic interactions favor the flux of viruses between phylogenetically distant organisms, such as insects and vertebrate predators. However, it is not yet possible to confirm viral infection at all trophic levels. The evolutionary tendency is for prey viruses to infect predators at some point, through frequent contact via feeding. While they are unable to infect (if this is the case, for example) by being ingested with their primary hosts, yet the predators serve as mechanical vectors that help disperse the viruses elsewhere, favoring horizontal transmission by the oral-fecal route [134] (especially if they are migratory animals such as birds and bats). Taking the example of the ALPV, the most prevalent and abundant virus in this analysis, found in four species of predators from three countries, even without a search focused on it, there is experimental evidence of its presence in the gut, lung, and spleen of two bat species (*Miniopterus schreibersii* and *Myotis chinensis*), see supplementary materials in [172]. In addition, the authors of that experimental study reported the presence of a high number of other viruses previously considered to be invertebrate-specific in species of wild small mammals. They further proposed that these mammals can be *bona fide* hosts for those viruses [135], which is consistent with the dispersal/transmission model featured here (Figure 7B). In addition, the ALPV and RhPV were detected in shrew lungs from China, as well as two other dicistroviruses [125]. Another study on bats from the Brazilian Amazon also detected the presence of one *Cripavirus* sp. in the lung [136]. One more *Cripavirus* sp. was detected in the blood of fruit bats and validated by RT-PCR [137]. In 2014, one dicistrovirus was detected in the lung of a French bat and it was validated by PCR [138]. A more recent research with lungs of shrews, hedgehogs, and shrew moles from China also reported the presence of dicistroviruses in its results [139]. A recent bat metagenomics compendium work revealed that arthropod and plant viruses flow into bats via diets, and insect-killing iflaviruses and dicistroviruses are common in the bat virome [140], which highlights that these flying mammals are good at spreading useful viruses in pest control [140]. This is highly relevant because the vast majority of virome studies on birds and small mammals focus on the detection of zoonotic viruses. Thus, these studies are anthropocentric and lack interpretations regarding viruses of agricultural and environmental importance occasionally detected. As a result, it is not yet possible to confirm whether viruses originally detected in plants and other arthropods are actually capable of infecting mammals. Because this is not an anthropocentric study, the principal message here is the possibility of viral dispersal through different levels, which can help the virus to reach different habitats, increasing the possibilities of infection on a large scale. This study is concerned with the conservation of beneficial insects, and improvement of biocontrol strategies, therefore it is not the aim to speculate on the replication of insect or plant viruses in the quoted vertebrate predators, although this hypothesis cannot be excluded.

Furthermore, cannibalism and/or intraguild predation among coexisting aphid-consuming insects [84] may contribute to horizontal transmission of microorganisms. Intraguild predation by ladybirds has been observed, primarily involving the consumption of eggs and larvae from other ladybirds [141].This may explain the presence of Harmonia axyridis virus 1, for instance, in other predators (ladybird and lacewing). Moreover, the evolutionary proximity of the new viruses found in organisms from different trophic levels (see Figure 5) suggests the possibility of cross-species viral transmission through the host’s diet. Recently, it has been shown that parasitoid wasp species share exogenous viruses with their prey and these viruses probably play a role in the success of the parasitoid-host relationship [142]. In addition, as reviewed by Islam et al. 2020, it is known that interactions involving plants, insect vectors, and viruses influence each other for their own survival, leading to the co-evolution among those involved. Also, other external abiotic and biotic players may influence these interactions [143].

### 4.5 Limitations on this study, but a world of testable possibilities

This data analysis produced interesting results, but with some limitations. The advantages of this study are its reproducibility, speed, low cost and wide geographical scope. However, it was not possible to carry out experiments aimed at the questions that arose from the insights gained. Still, the results obtained point towards more applied studies to solve problems in agriculture and insect conservation. It is truly astonishing that insect species are commercialized and released in various regions without at least one study on their viruses. For these reasons, this work does not intend to solve all the questions about the viruses, their hosts, transmission routes and ecological interactions, but to show that there is much to be investigated in this area. Therefore, the findings here reported may inspire studies following the traditional scientific method – hypothesis-based deduction, which implies that a falsifiable hypothesis needs to be tested experimentally or computationally, and the results should support or refute the hypothesis, triggering a new round of formulation and testing [144].

In conclusion, the simple assessment of viral diversity in eight predatory insects of agricultural relevance using a bioinformatics approach has revealed a possible interspecies viral flux that deserves research attention. Hence, it was suggested that interactions involving plants, insect vector, viruses, natural enemies, and other predators (at trophic levels above) may influence or be influenced by each other, in the context of an agroecological system. Therefore, as often as possible, a broader approach to virus surveillance should be considered when studying predator species, such as biological control agents. This is due to the fact that a community or ecosystem perspective can provide more evolutionary and ecological insights on viruses than studying single host species. For this reason, understanding the role of viruses in trophic relationships of economic importance requires further research. Consequently, the development of enhanced biological control, agricultural productivity, sustainability, and conservation strategies may be achieved.

## 5. Data availability statement

The public RNA-seq libraries analyzed in this study are available in the SRA - NCBI repository. The nucleotide sequence data reported in this work are available from the Third-Party Annotation Section of the DDBJ/ENA/GenBank databases under the accession numbers TPA: BK071453-BK071525.

## 6. Author contributions

G.B.C-G.: Conceptualization, Methodology, Formal analysis and investigation, Writing – original draft preparation, Writing – review and editing, and Visualization; I.S.L.: Methodology and Visualization; E.R.G.R.A.: Conceptualization, Methodology, Writing – review and editing, Visualization, Resources, and Supervision. All authors read and approved the manuscript.

## Acknowledgments

We thank all the scientists at the Virus Bioinformatics Laboratory–UESC for the meaningful discussions, the Center of Biotechnology and Genetics, and LabiOmicas for providing the infrastructure to carry out the research.

