## Supplementary text for "The Initial Detection of a Diversity of Viruses Associated with Ladybirds and Other Biocontrol Agents Prompts Interesting Ecological Insights"

On the diversity of new virus discoveries

The 41 new viruses were classified into 16 families and 1 genus (**Figure 2B**). New viruses of *Iflaviridae* and *Partitiviridae* were found in the three groups (ladybirds, lacewings, and aphid midges). Iflaviruses have ssRNA(+) genomes of 9-11 kb in size. They are arthropod viruses, mostly infecting insects. The main route of infection is the ingestion of contaminated food. Beneficial and harmful insects host iflaviruses that can be asymptomatic or cause illness (developmental abnormalities, behavioral changes, and premature mortality) [1]. Partitiviruses have segmented dsRNA genomes of 3-4.8 kb [2]. Although at first associated with plant or fungal hosts [2], several partitiviruses have been discovered in insects in recent years [3–6]. In addition, new *Dicistroviridae* members were found in ladybirds (**Figure 1B**). With ssRNA(+) genomes of 8-10kb, dicistroviruses are common viruses of arthropods [7], some with pathogenic effects on bees [8]. Also, dicistroviruses have been found in mammals (gorillas, giant pandas, squirrel, and bats) [9–13]. Ladybirds also had phasmaviruses, ssRNA(-) viruses of 9.7-15.8 kb in size that are maintained in insects [14]. *Lispiviridae* members were found in a ladybird and a lacewing. Lispiviruses have ssRNA(-) genomes of 6.5-15.5 kb, and have been found in arthropods and nematodes [15]. Picornaviruses or picorna-like viruses were found in ladybirds and lacewings. Members of *Picornaviridae* have ssRNA(+) genomes of 6.7 to 10.1 kb and infect vertebrate hosts, such as mammals, birds, reptiles, amphibians, and fish [16]. However, 'picorna-like' viruses have been found in several arthropod hosts, such as mosquitoes [17,18], mites [19], aphids [20], and a ladybird [21]. A *Secoviridae* new member was found in a ladybird (**Figure 2B**). Secoviruses can have segmented or non-segmented ssRNA(+) genomes of 9-13.7 kb in size [22]. Although at first associated with plants, secoviruses have been found in bees [23], ticks [24], among others [25]. Furthermore, a new polycipivirus was detected in a ladybird. Polycipiviruses have polycistronic ssRNA(+) genomes of 10-12kb in size [26] that have been found in insects

[4,27] and birds [28]. A member of *Solinviviridae* was detected in a ladybird. Solinviviruses have ssRNA(+) genomes of 10-11kb, and they have been found in insects and other arthropods [29]. Partial spinareoviruses were found in a ladybird and a lacewing (**Figure 2B**). These viruses have segmented dsRNA genomes of 9-12 segments totalizing 23-29 kbp in size [30]. These viruses infect a wide range of hosts, including animals, fungi, and plants, with some exhibiting pathogenicity [30]. Amalgaviruses have dsRNA genomes and infect plants, fungi, and animals [31]. An amalgavirus was detected in a lacewing (**Figure 2B**). Further, narnaviruses also were detected in lacewings. These viruses have ssRNA(+) genomes of 2.3-3.6 kb in size [32], and have been found in fungi [33] and insects [27,34]. Negeviruses were found in a lacewing and in the aphid midge (**Figure 2B**). These viruses have ssRNA(+) genomes with poly(A) tails, and have been found in hematophagous insects (mosquitoes and sand flies) [35,36]. A member of *Orthototiviridae* was detected in a lacewing. Orthototiviruses have a dsRNA genome of 3.9-5.6 kb in size, and have been detected in plants, protists, fungi, and invertebrates [37,38]. In addition, a rhabdovirus was found in a lacewing. Members of *Rhabdoviridae* have ssRNA(-) of 10-16 kb and infect plants or animals (invertebrates and vertebrates). Rhabdoviruses can be transmitted by arthropods and may cause disease in mammals, fish, and farm crops [39]. Also, a solemovirus was detected in a lacewing. Members of *Solemoviridae* have polycistronic ssRNA(+) genomes of 4-6 kb in size, and they infect plants. Some solemoviruses are transmitted by aphids and have high pathogenicity and economic impact [40]. Tymoviruses were identified in a lacewing and in the aphid midge (**Figure 2B**). *Tymoviridae* members have ssRNA(+) genomes of 6.0-7.5 kb in size and infect plants and animals [41].
