## Supplementary material for "The Initial Detection of a Diversity of Viruses Associated with Ladybirds and Other Biocontrol Agents Prompts Interesting Ecological Insights"

### *Amalgaviridae*

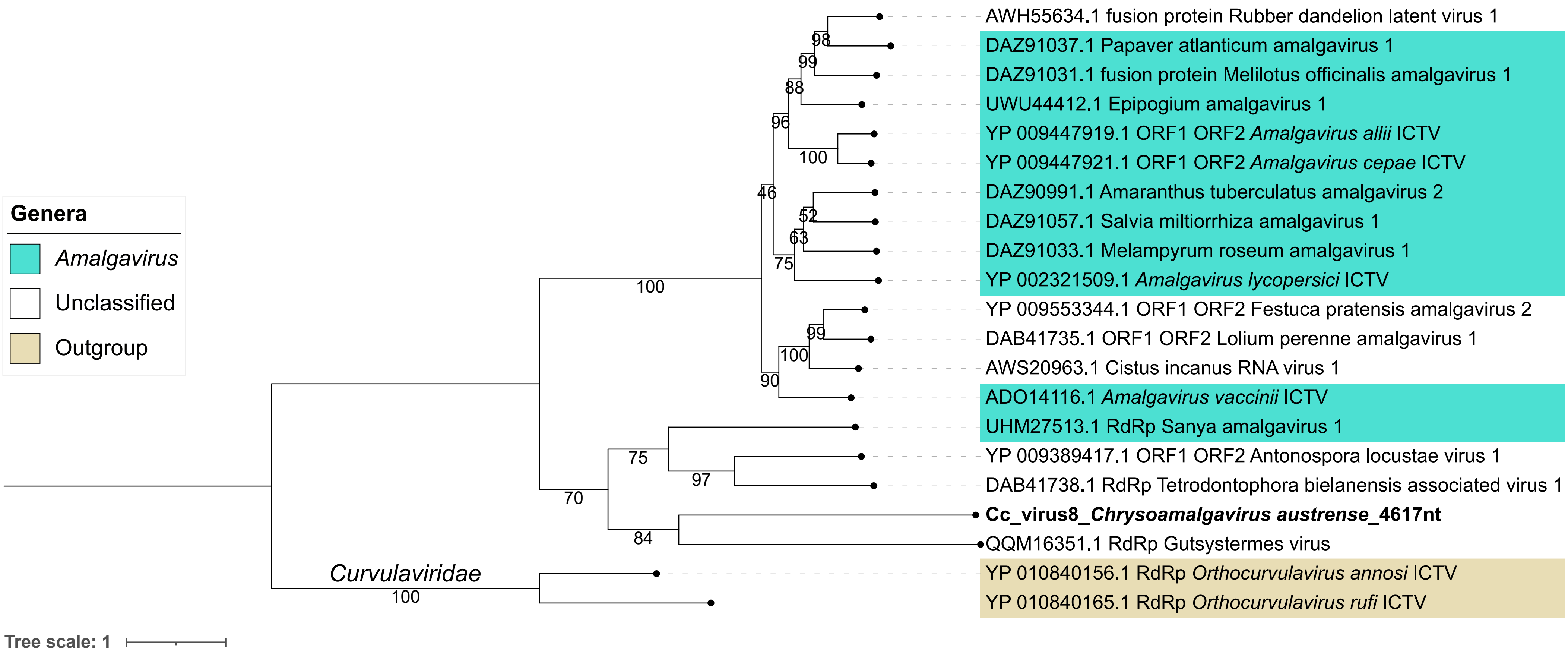

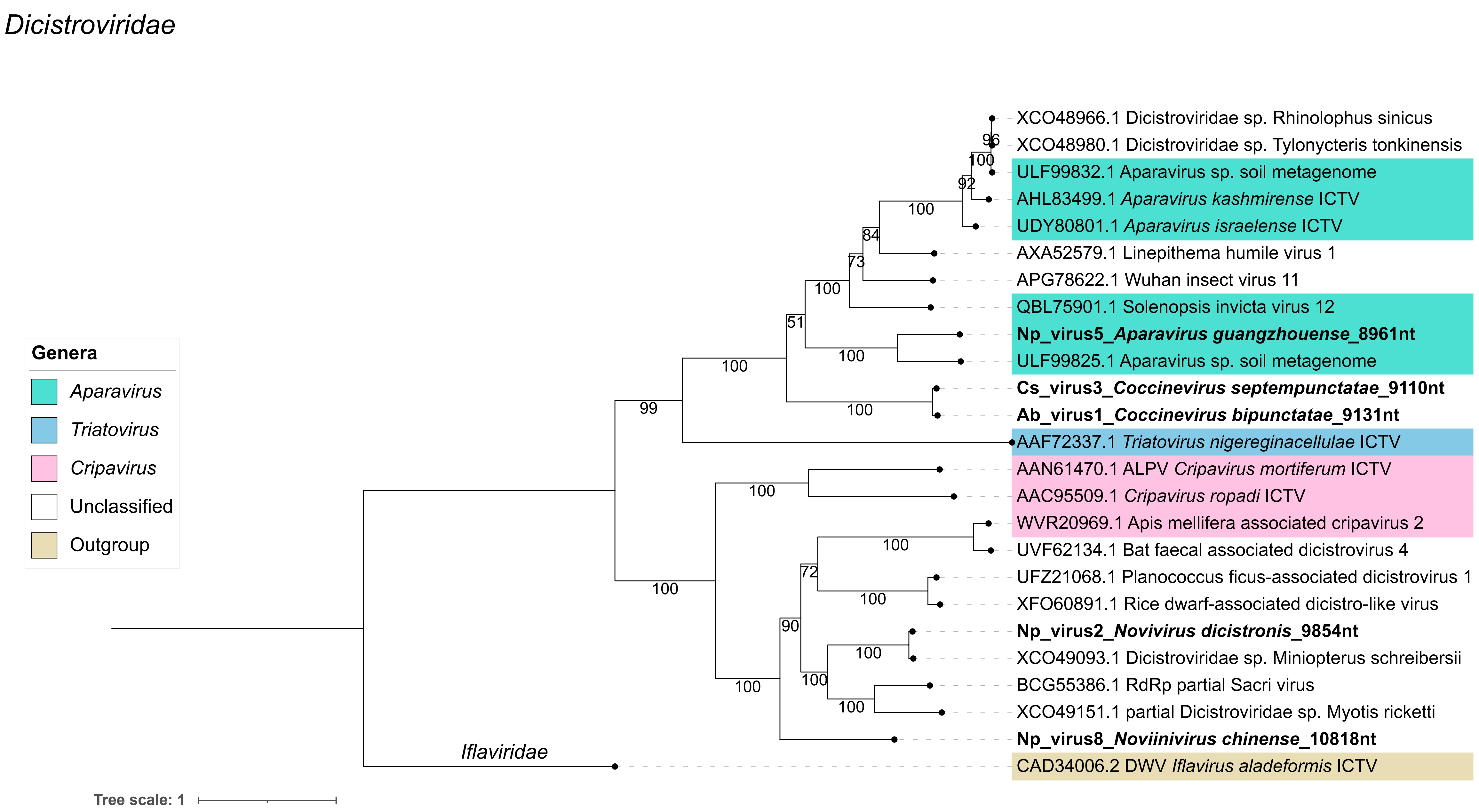
Dicistroviridae
Genera
Aparavirus
Triatovirus
Cripavirus
Unclassified
Outgroup
XCO48966.1 Dicistroviridae sp. Rhinolophus sinicus
XCO48980.1 Dicistroviridae sp. Tylonycteris tonkinensis
ULF99832.1 Aparavirus sp. soil metagenome
AHL83499.1 Aparavirus kashmirense ICTV
UDY80801.1 Aparavirus israelense ICTV
AXA52579.1 Linepithema humile virus 1
APG78622.1 Wuhan insect virus 11
QBL75901.1 Solenopsis invicta virus 12
Np_virus5_Aparavirus guangzhouense_8961nt
ULF99825.1 Aparavirus sp. soil metagenome
Cs_virus3_Coccinevirus septempunctatae_9110nt
Ab_virus1_Coccinevirus bipunctatae_9131nt
AAF72337.1 Triatovirus nigereginacellulae ICTV
AAN61470.1 ALPV Cripavirus mortiferum ICTV
AAC95509.1 Cripavirus ropadi ICTV
WVR20969.1 Apis mellifera associated cripavirus 2
UVF62134.1 Bat faecal associated dicistrovirus 4
UFZ21068.1 Planococcus ficus-associated dicistrovirus 1
XFO60891.1 Rice dwarf-associated dicistro-like virus
Np_virus2_Novivirus dicistronis_9854nt
XCO49093.1 Dicistroviridae sp. Miniopterus schreibersii
BCG55386.1 RdRp partial Sacri virus
XCO49151.1 partial Dicistroviridae sp. Myotis ricketti
Np_virus8_Noviinivirus chinense_10818nt
CAD34006.2 DWV Iflavirus aladeformis ICTV
Iflaviridae
Tree scale: 1
96
100
92
100
84
73
100
51
100
100
100
99
100
100
100
72
100
90
100
100
100
100

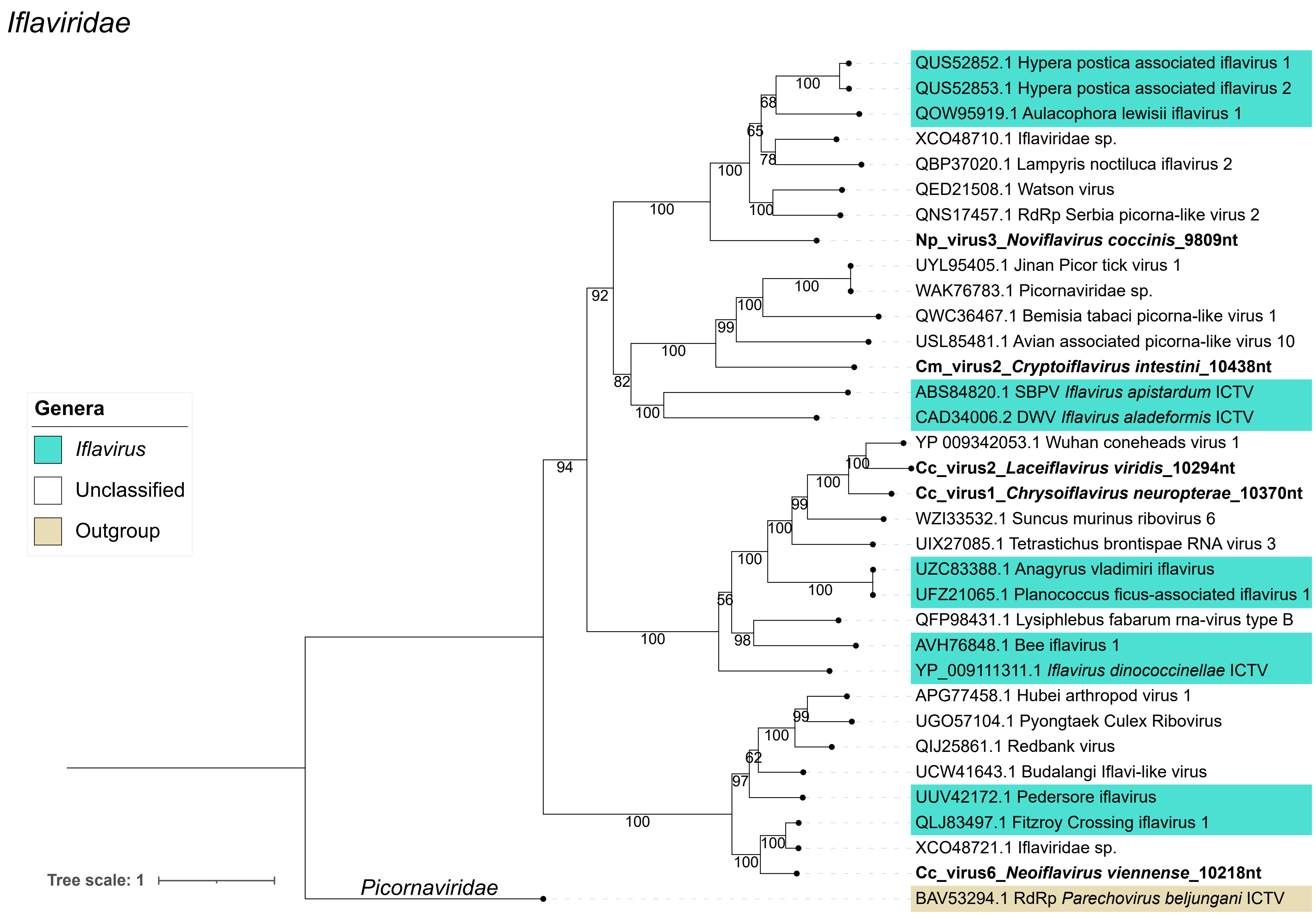

Iflaviridae
QUS52852.1 Hypera postica associated iflavirus 1
QUS52853.1 Hypera postica associated iflavirus 2
QOW95919.1 Aulacophora lewisii iflavirus 1
XCO48710.1 Iflaviridae sp.
QBP37020.1 Lampyris noctiluca iflavirus 2
QED21508.1 Watson virus
QNS17457.1 RdRp Serbia picorna-like virus 2
Np_virus3_Noviflavirus coccinis_9809nt
UYL95405.1 Jinan Picor tick virus 1
WAK76783.1 Picornaviridae sp.
QWC36467.1 Bemisia tabaci picorna-like virus 1
USL85481.1 Avian associated picorna-like virus 10
Cm_virus2_Cryptoiflavirus intestini_10438nt
ABS84820.1 SBPV Iflavirus apistardum ICTV
CAD34006.2 DWV Iflavirus aladeformis ICTV
YP 009342053.1 Wuhan coneheads virus 1
Cc_virus2_Laceiflavirus viridis_10294nt
Cc_virus1_Chrysoiflavirus neuropterae_10370nt
WZI33532.1 Suncus murinus ribovirus 6
UIX27085.1 Tetrastichus brontispae RNA virus 3
UZC83388.1 Anagyrus vladimiri iflavirus
UFZ21065.1 Planococcus ficus-associated iflavirus 1
QFP98431.1 Lysiphlebus fabarum rna-virus type B
AVH76848.1 Bee iflavirus 1
YP_009111311.1 Iflavirus dinococcinellae ICTV
APG77458.1 Hubei arthropod virus 1
UGO57104.1 Pyongtaek Culex Ribovirus
QIJ25861.1 Redbank virus
UCW41643.1 Budalangi Iflavi-like virus
UUV42172.1 Pedersore iflavirus
QLJ83497.1 Fitzroy Crossing iflavirus 1
XCO48721.1 Iflaviridae sp.
Cc_virus6_Neoiflavirus viennense_10218nt
BAV53294.1 RdRp Parechovirus beljungani ICTV
Picornaviridae
Genera
Iflavirus
Unclassified
Outgroup
Tree scale: 1
100
68
65
78
100
100
100
92
100
100
99
100
82
100
94
100
100
99
100
100
100
56
98
100
99
100
62
97
100
100
100

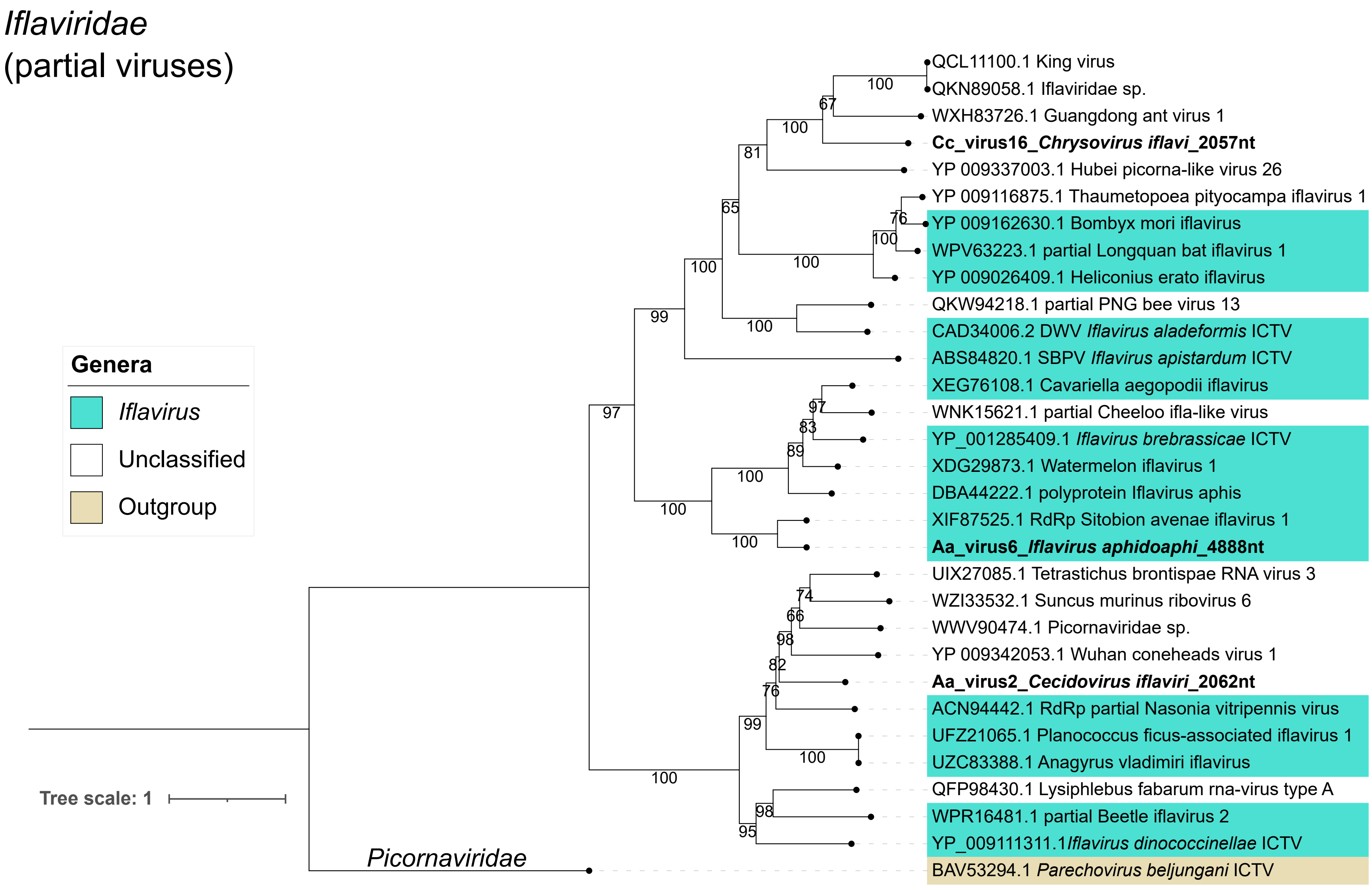

Iflaviridae
(partial viruses)
Genera
Iflavirus
Unclassified
Outgroup
Tree scale: 1
Picornaviridae
QCL11100.1 King virus
QKN89058.1 Iflaviridae sp.
WXH83726.1 Guangdong ant virus 1
Cc_virus16_Chrysovirus iflavi_2057nt
YP 009337003.1 Hubei picorna-like virus 26
YP 009116875.1 Thaumetopoea pityocampa iflavirus 1
YP 009162630.1 Bombyx mori iflavirus
WPV63223.1 partial Longquan bat iflavirus 1
YP 009026409.1 Heliconius erato iflavirus
QKW94218.1 partial PNG bee virus 13
CAD34006.2 DWV Iflavirus aladeformis ICTV
ABS84820.1 SBPV Iflavirus apistardum ICTV
XEG76108.1 Cavariella aegopodii iflavirus
WNK15621.1 partial Cheeloo ifla-like virus
YP_001285409.1 Iflavirus brebrassicae ICTV
XDG29873.1 Watermelon iflavirus 1
DBA44222.1 polyprotein Iflavirus aphis
XIF87525.1 RdRp Sitobion avenae iflavirus 1
Aa_virus6_Iflavirus aphidoaphi_4888nt
UIX27085.1 Tetrastichus brontispae RNA virus 3
WZI33532.1 Suncus murinus ribovirus 6
WWV90474.1 Picornaviridae sp.
YP 009342053.1 Wuhan coneheads virus 1
Aa_virus2_Cecidovirus iflaviri_2062nt
ACN94442.1 RdRp partial Nasonia vitripennis virus
UFZ21065.1 Planococcus ficus-associated iflavirus 1
UZC83388.1 Anagyrus vladimiri iflavirus
QFP98430.1 Lysiphlebus fabarum rna-virus type A
WPR16481.1 partial Beetle iflavirus 2
YP_009111311.1Iflavirus dinococcinellae ICTV
BAV53294.1 Parechovirus beljungani ICTV

*Lispiviridae*

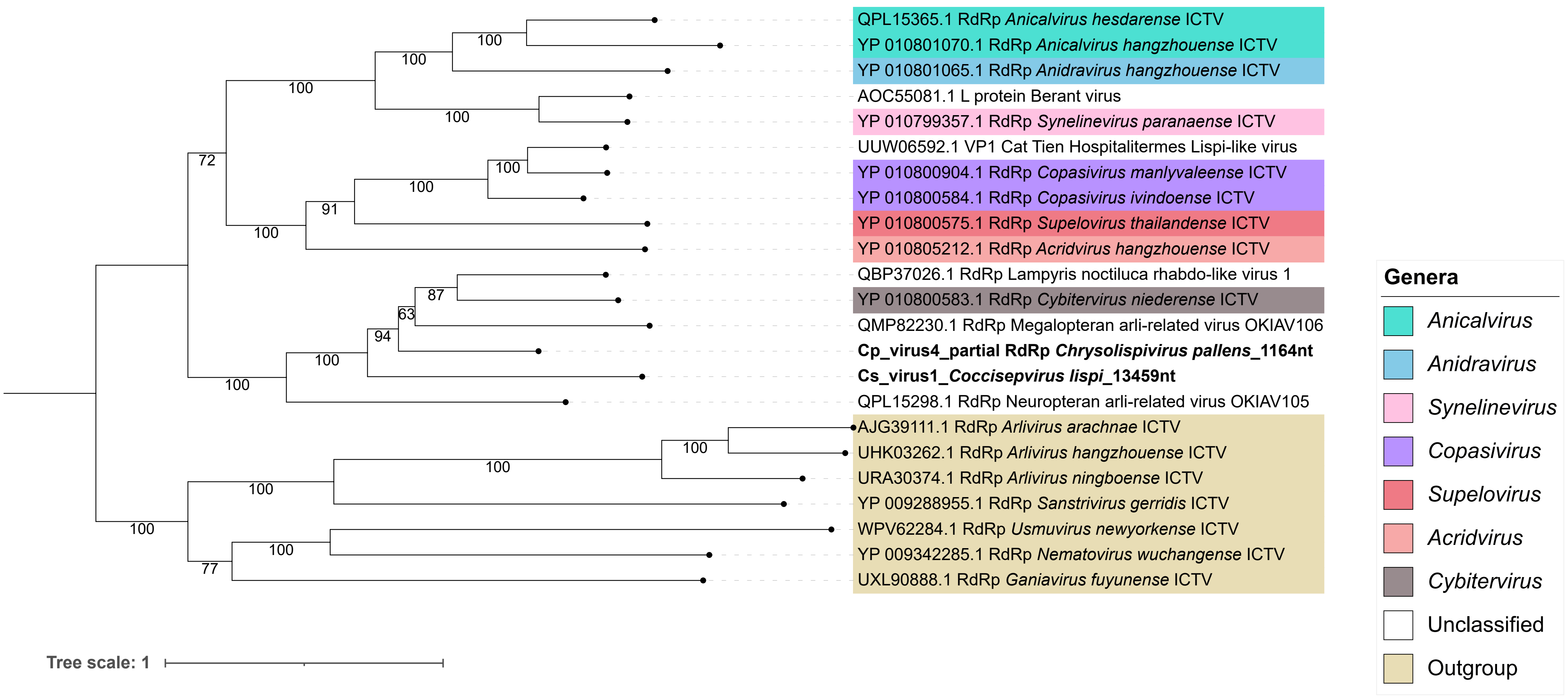

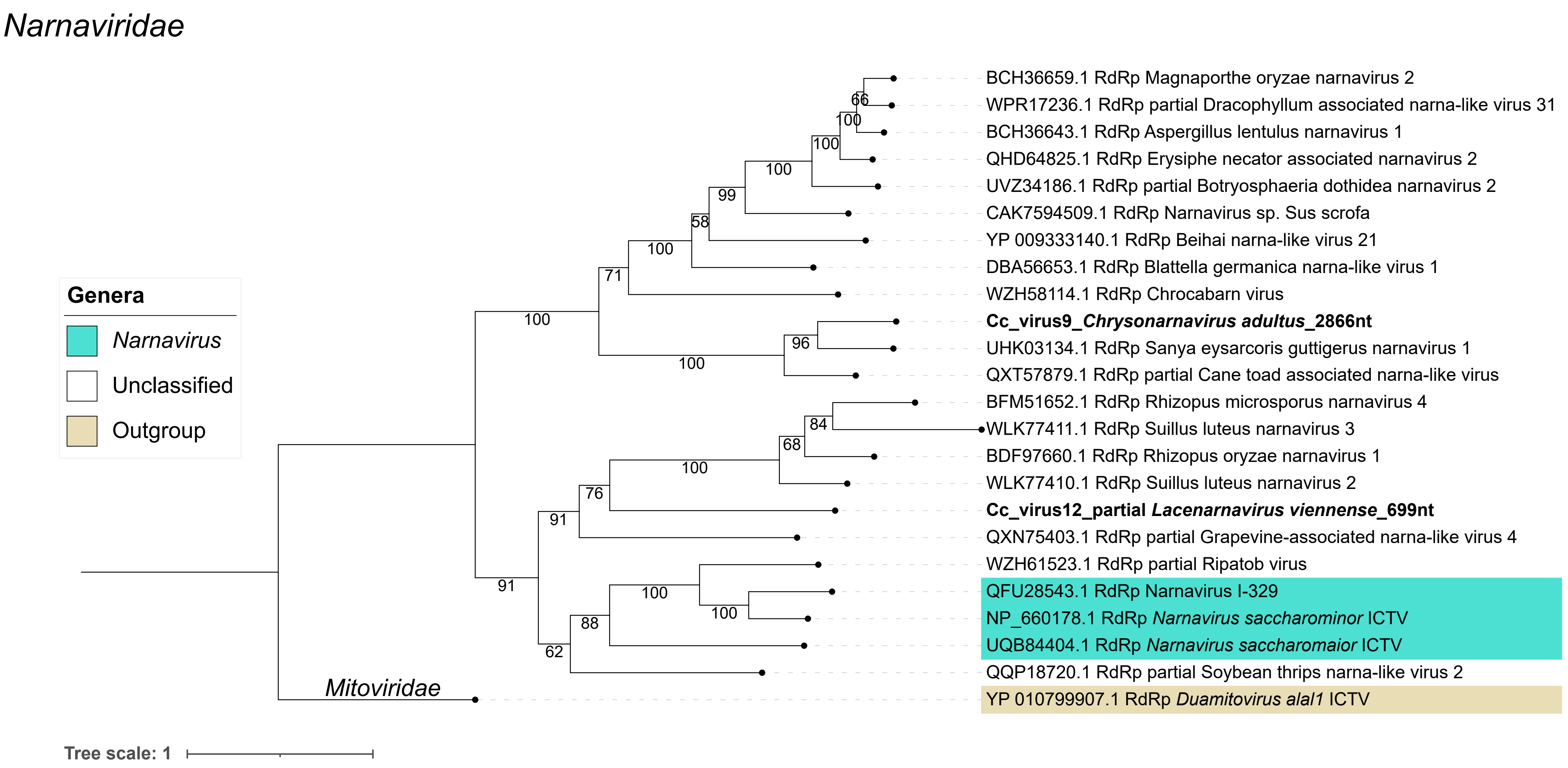

Narnaviridae
Genera
Narnavirus
Unclassified
Outgroup
BCH36659.1 RdRp Magnaporthe oryzae narnavirus 2
WPR17236.1 RdRp partial Dracophyllum associated narna-like virus 31
BCH36643.1 RdRp Aspergillus lentulus narnavirus 1
QHD64825.1 RdRp Erysiphe necator associated narnavirus 2
UVZ34186.1 RdRp partial Botryosphaeria dothidea narnavirus 2
CAK7594509.1 RdRp Narnavirus sp. Sus scrofa
YP 009333140.1 RdRp Beihai narna-like virus 21
DBA56653.1 RdRp Blattella germanica narna-like virus 1
WZH58114.1 RdRp Chrocabarn virus
Cc_virus9_Chrysonarnavirus adultus_2866nt
UHK03134.1 RdRp Sanya eysarcoris guttigerus narnavirus 1
QXT57879.1 RdRp partial Cane toad associated narna-like virus
BFM51652.1 RdRp Rhizopus microsporus narnavirus 4
WLK77411.1 RdRp Suillus luteus narnavirus 3
BDF97660.1 RdRp Rhizopus oryzae narnavirus 1
WLK77410.1 RdRp Suillus luteus narnavirus 2
Cc_virus12_partial Lacenarnavirus viennense_699nt
QXN75403.1 RdRp partial Grapevine-associated narna-like virus 4
WZH61523.1 RdRp partial Ripatob virus
QFU28543.1 RdRp Narnavirus I-329
NP_660178.1 RdRp Narnavirus saccharominor ICTV
UQB84404.1 RdRp Narnavirus saccharomaior ICTV
QQP18720.1 RdRp partial Soybean thrips narna-like virus 2
YP 010799907.1 RdRp Duamitovirus alal1 ICTV
Mitoviridae
Tree scale: 1

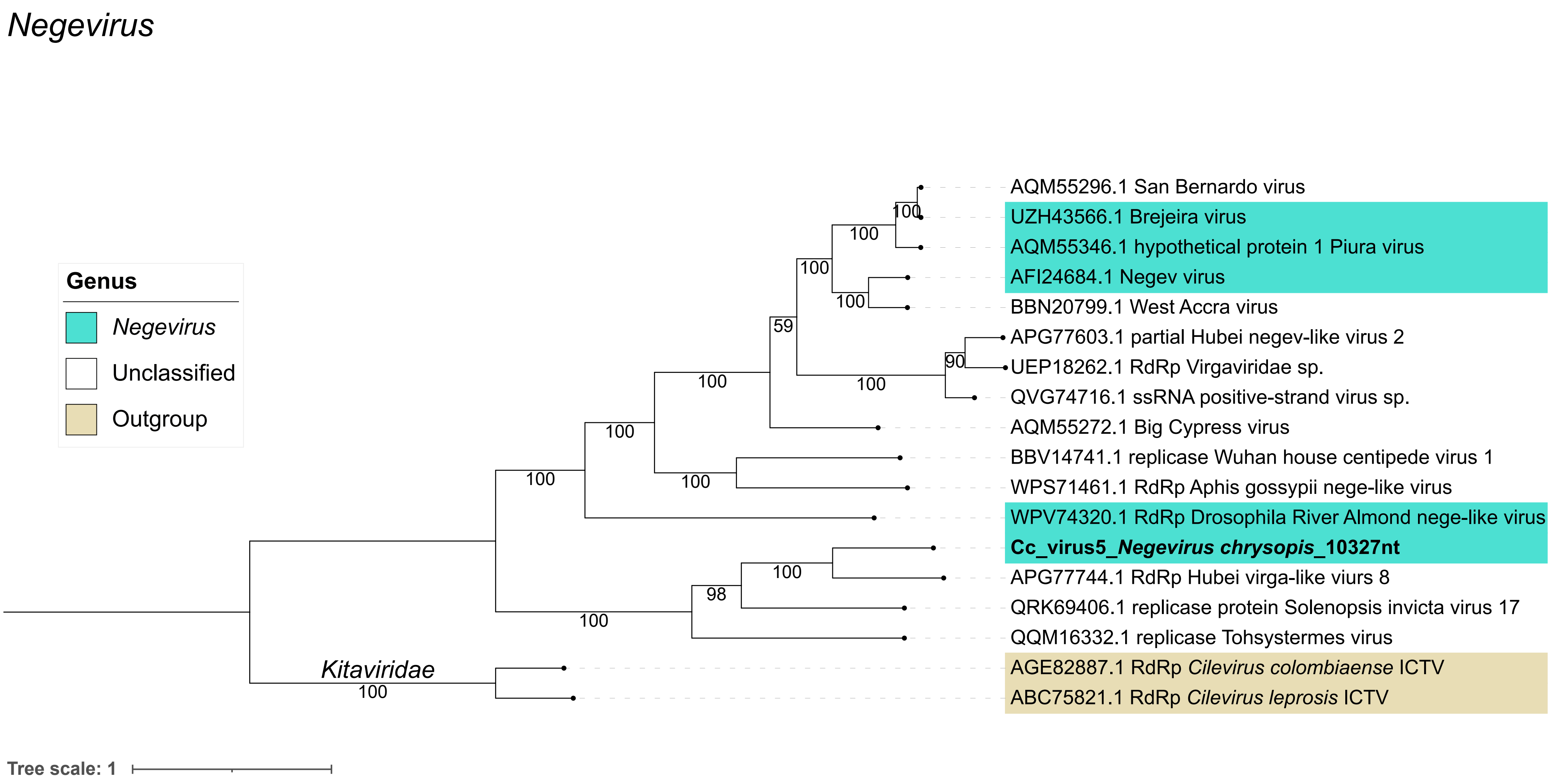
Negevirus
Genus
Negevirus
Unclassified
Outgroup
AQM55296.1 San Bernardo virus
UZH43566.1 Brejeira virus
AQM55346.1 hypothetical protein 1 Piura virus
AFI24684.1 Negev virus
BBN20799.1 West Accra virus
APG77603.1 partial Hubei negev-like virus 2
UEP18262.1 RdRp Virgaviridae sp.
QVG74716.1 ssRNA positive-strand virus sp.
AQM55272.1 Big Cypress virus
BBV14741.1 replicase Wuhan house centipede virus 1
WPS71461.1 RdRp Aphis gossypii nege-like virus
WPV74320.1 RdRp Drosophila River Almond nege-like virus
Cc_virus5_Negevirus chrysopis_10327nt
APG77744.1 RdRp Hubei virga-like viurs 8
QRK69406.1 replicase protein Solenopsis invicta virus 17
QQM16332.1 replicase Tohsystermes virus
AGE82887.1 RdRp Cilevirus colombiaense ICTV
ABC75821.1 RdRp Cilevirus leprosis ICTV
Kitaviridae
Tree scale: 1

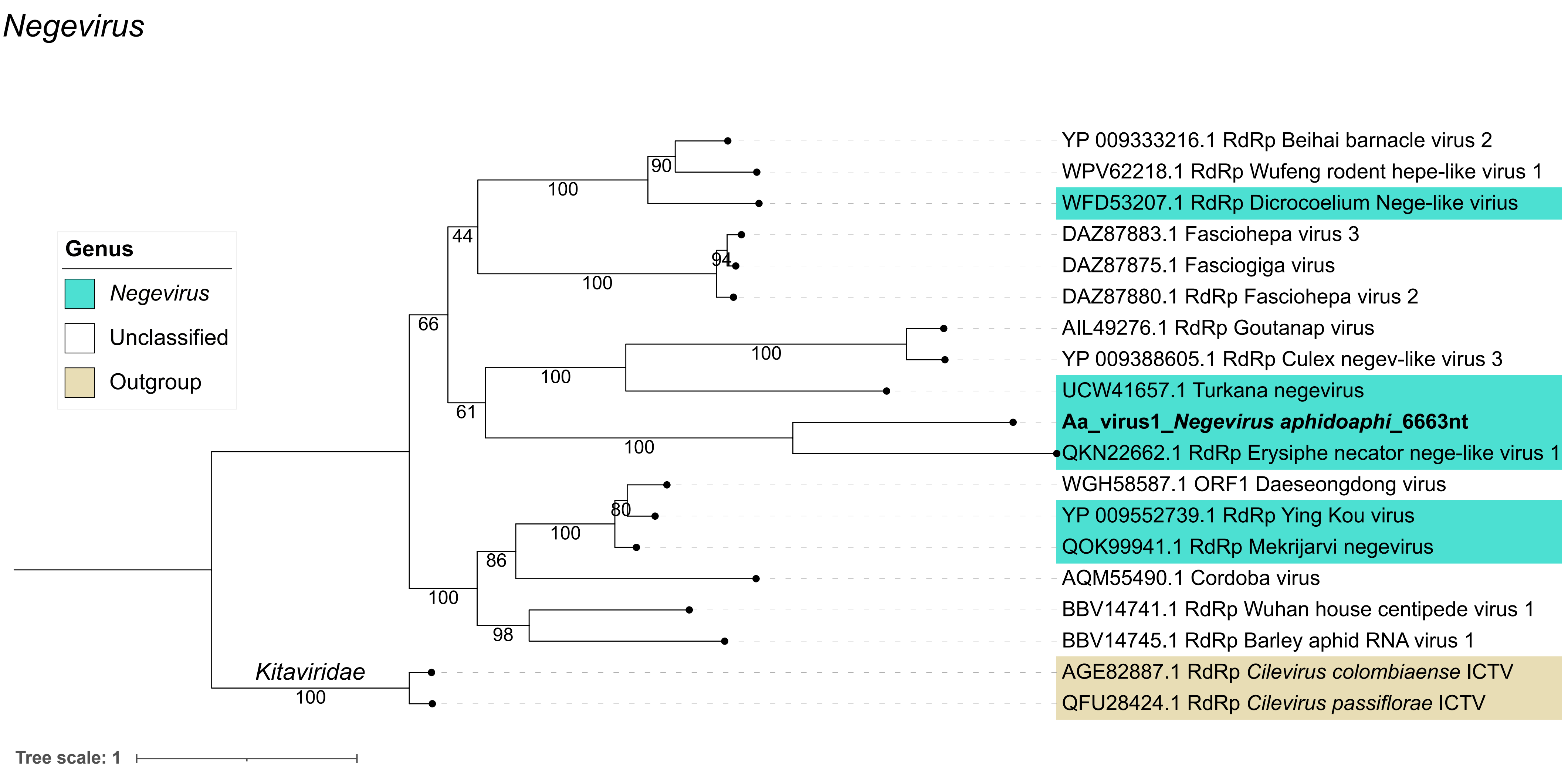
Negevirus
Genus
Negevirus
Unclassified
Outgroup
YP 009333216.1 RdRp Beihai barnacle virus 2
WPV62218.1 RdRp Wufeng rodent hepe-like virus 1
WFD53207.1 RdRp Dicrocoelium Nege-like virius
DAZ87883.1 Fasciohepa virus 3
DAZ87875.1 Fasciogiga virus
DAZ87880.1 RdRp Fasciohepa virus 2
AIL49276.1 RdRp Goutanap virus
YP 009388605.1 RdRp Culex negev-like virus 3
UCW41657.1 Turkana negevirus
Aa_virus1_Negevirus aphidoaphi_6663nt
QKN22662.1 RdRp Erysiphe necator nege-like virus 1
WGH58587.1 ORF1 Daeseongdong virus
YP 009552739.1 RdRp Ying Kou virus
QOK99941.1 RdRp Mekrijarvi negevirus
AQM55490.1 Cordoba virus
BBV14741.1 RdRp Wuhan house centipede virus 1
BBV14745.1 RdRp Barley aphid RNA virus 1
AGE82887.1 RdRp Cilevirus colombiaense ICTV
QFU28424.1 RdRp Cilevirus passiflorae ICTV
Kitaviridae
90
100
44
94
100
66
100
100
61
100
80
100
86
100
98
100
Tree scale: 1

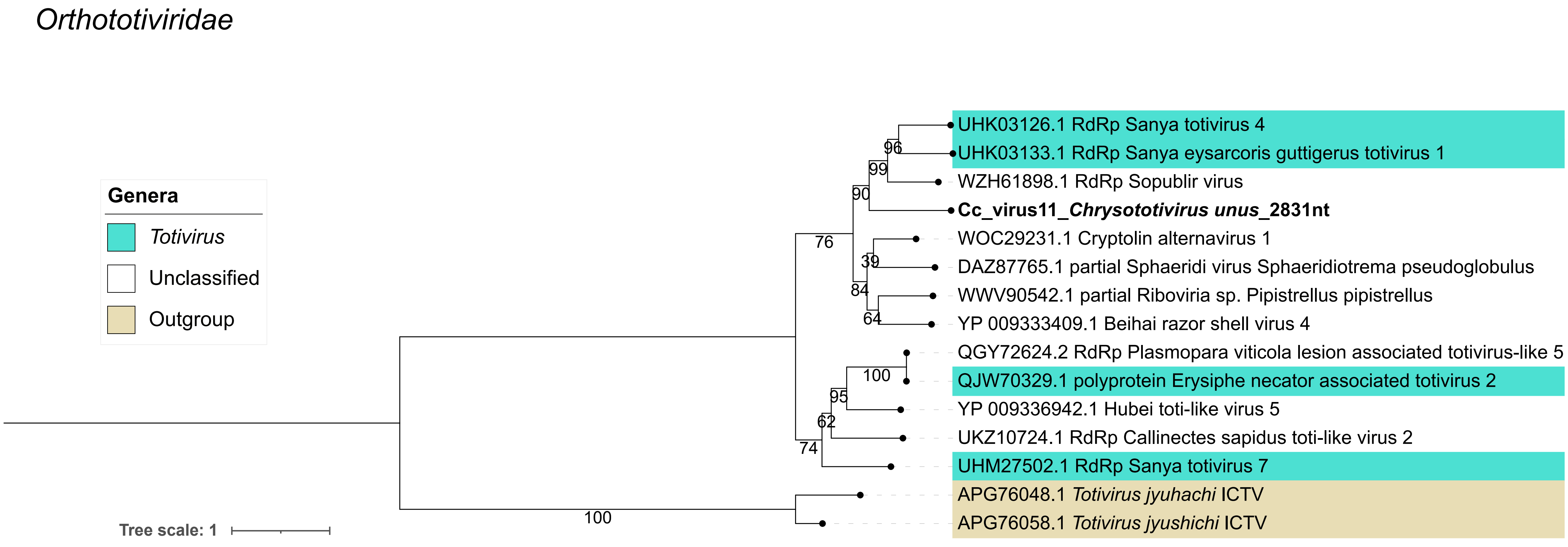

Orthototiviridae
Genera
Totivirus
Unclassified
Outgroup
Tree scale: 1
UHK03126.1 RdRp Sanya totivirus 4
UHK03133.1 RdRp Sanya eysarcoris guttigerus totivirus 1
WZH61898.1 RdRp Sopublir virus
Cc_virus11_Chrysototivirus unus_2831nt
WOC29231.1 Cryptolin alternavirus 1
DAZ87765.1 partial Sphaeridi virus Sphaeridiotrema pseudoglobulus
WWV90542.1 partial Riboviria sp. Pipistrellus pipistrellus
YP 009333409.1 Beihai razor shell virus 4
QGY72624.2 RdRp Plasmopara viticola lesion associated totivirus-like 5
QJW70329.1 polyprotein Erysiphe necator associated totivirus 2
YP 009336942.1 Hubei toti-like virus 5
UKZ10724.1 RdRp Callinectes sapidus toti-like virus 2
UHM27502.1 RdRp Sanya totivirus 7
APG76048.1 Totivirus jyuhachi ICTV
APG76058.1 Totivirus jyushichi ICTV
96
99
90
76
39
84
64
100
95
62
74
100

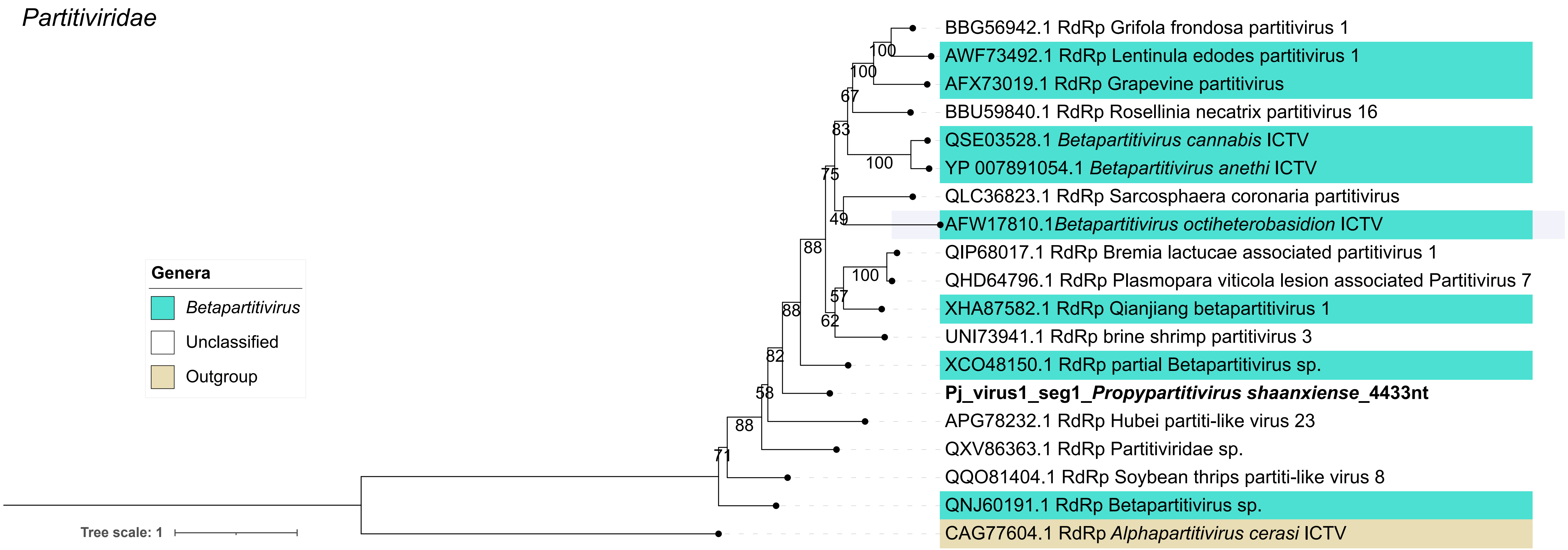

Partitiviridae
BBG56942.1 RdRp Grifola frondosa partitivirus 1
AWF73492.1 RdRp Lentinula edodes partitivirus 1
AFX73019.1 RdRp Grapevine partitivirus
BBU59840.1 RdRp Rosellinia necatrix partitivirus 16
QSE03528.1 Betapartitivirus cannabis ICTV
YP 007891054.1 Betapartitivirus anethi ICTV
QLC36823.1 RdRp Sarcosphaera coronaria partitivirus
AFW17810.1Betapartitivirus octiheterobasidion ICTV
QIP68017.1 RdRp Bremia lactucae associated partitivirus 1
QHD64796.1 RdRp Plasmopara viticola lesion associated Partitivirus 7
XHA87582.1 RdRp Qianjiang betapartitivirus 1
UNI73941.1 RdRp brine shrimp partitivirus 3
XCO48150.1 RdRp partial Betapartitivirus sp.
Pj_virus1_seg1_Propypartitivirus shaanxiense_4433nt
APG78232.1 RdRp Hubei partiti-like virus 23
QXV86363.1 RdRp Partitiviridae sp.
QQO81404.1 RdRp Soybean thrips partiti-like virus 8
QNJ60191.1 RdRp Betapartitivirus sp.
CAG77604.1 RdRp Alphapartitivirus cerasi ICTV
Genera
Betapartitivirus
Unclassified
Outgroup
Tree scale: 1
100
100
67
83
100
75
49
88
100
57
88
62
82
58
88
71

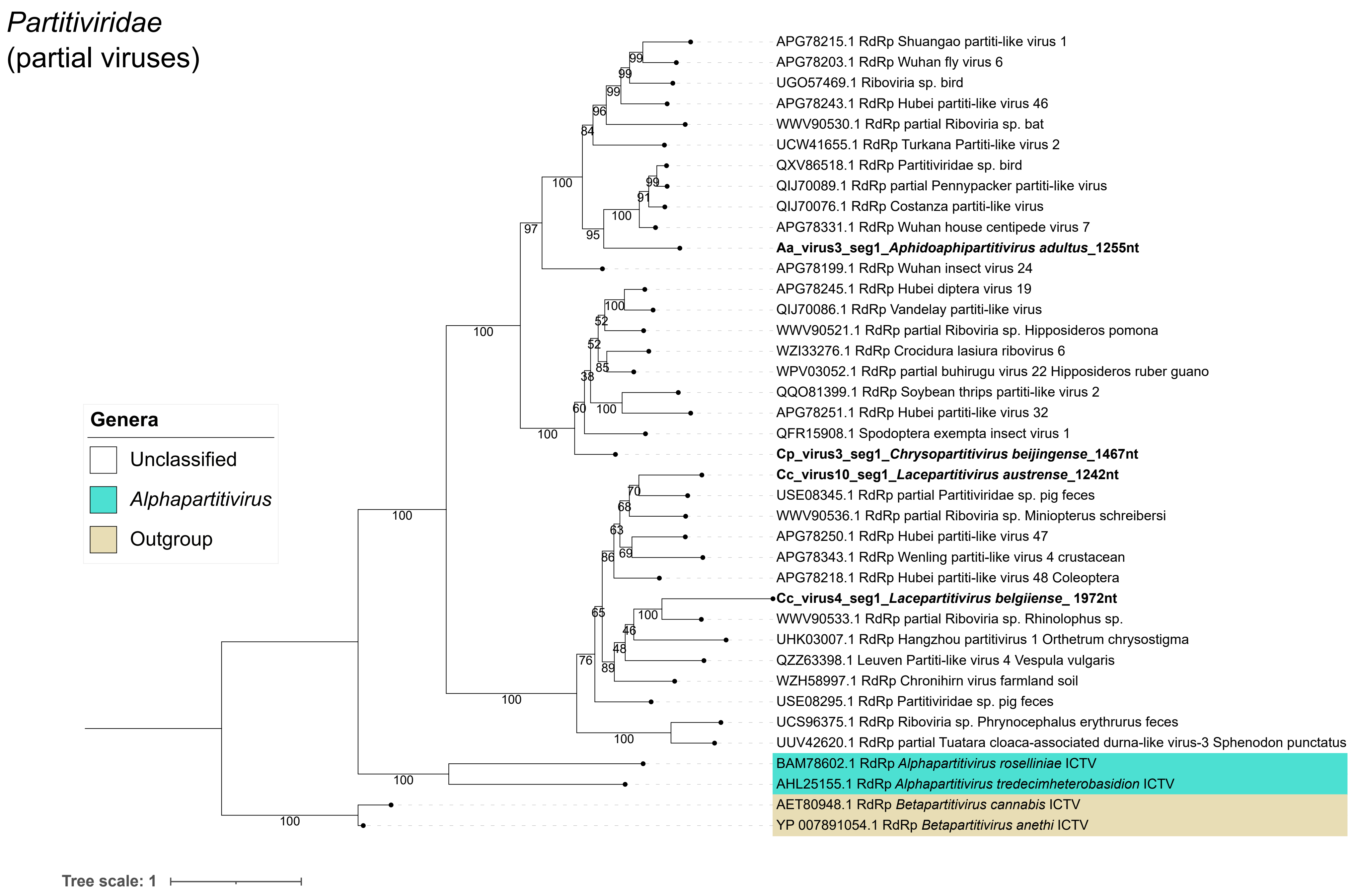
Partitiviridae
(partial viruses)
Genera
Unclassified
Alphapartitivirus
Outgroup
APG78215.1 RdRp Shuangao partiti-like virus 1
APG78203.1 RdRp Wuhan fly virus 6
UGO57469.1 Riboviria sp. bird
APG78243.1 RdRp Hubei partiti-like virus 46
WWV90530.1 RdRp partial Riboviria sp. bat
UCW41655.1 RdRp Turkana Partiti-like virus 2
QXV86518.1 RdRp Partitiviridae sp. bird
QIJ70089.1 RdRp partial Pennypacker partiti-like virus
QIJ70076.1 RdRp Costanza partiti-like virus
APG78331.1 RdRp Wuhan house centipede virus 7
Aa_virus3_seg1_Aphidoaphipartitivirus adultus_1255nt
APG78199.1 RdRp Wuhan insect virus 24
APG78245.1 RdRp Hubei diptera virus 19
QIJ70086.1 RdRp Vandelay partiti-like virus
WWV90521.1 RdRp partial Riboviria sp. Hipposideros pomona
WZI33276.1 RdRp Crocidura lasiura ribovirus 6
WPV03052.1 RdRp partial buhirugu virus 22 Hipposideros ruber guano
QQO81399.1 RdRp Soybean thrips partiti-like virus 2
APG78251.1 RdRp Hubei partiti-like virus 32
QFR15908.1 Spodoptera exempta insect virus 1
Cp_virus3_seg1_Chrysopartitivirus beijingense_1467nt
Cc_virus10_seg1_Lacepartitivirus austrense_1242nt
USE08345.1 RdRp partial Partitiviridae sp. pig feces
WWV90536.1 RdRp partial Riboviria sp. Miniopterus schreibersi
APG78250.1 RdRp Hubei partiti-like virus 47
APG78343.1 RdRp Wenling partiti-like virus 4 crustacean
APG78218.1 RdRp Hubei partiti-like virus 48 Coleoptera
Cc_virus4_seg1_Lacepartitivirus belgiiense_ 1972nt
WWV90533.1 RdRp partial Riboviria sp. Rhinolophus sp.
UHK03007.1 RdRp Hangzhou partitivirus 1 Orthetrum chrysostigma
QZZ63398.1 Leuven Partiti-like virus 4 Vespula vulgaris
WZH58997.1 RdRp Chronihirn virus farmland soil
USE08295.1 RdRp Partitiviridae sp. pig feces
UCS96375.1 RdRp Riboviria sp. Phrynocephalus erythrurus feces
UUV42620.1 RdRp partial Tuatara cloaca-associated durna-like virus-3 Sphenodon punctatus
BAM78602.1 RdRp Alphapartitivirus roselliniae ICTV
AHL25155.1 RdRp Alphapartitivirus tredecimheterobasidion ICTV
AET80948.1 RdRp Betapartitivirus cannabis ICTV
YP 007891054.1 RdRp Betapartitivirus anethi ICTV
Tree scale: 1

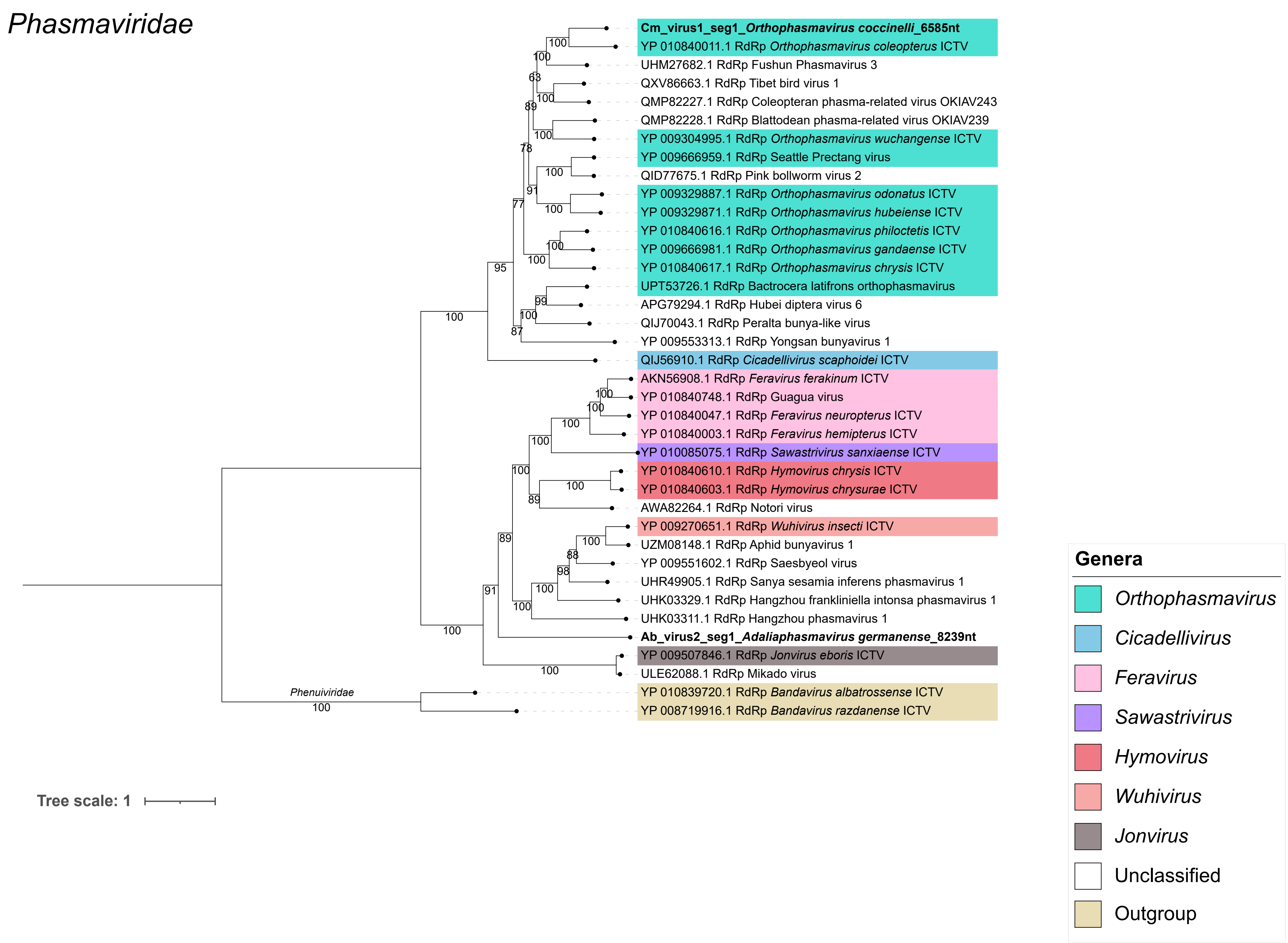

Phasmaviridae
Cm_virus1_seg1_Orthophasmavirus coccinelli_6585nt
YP 010840011.1 RdRp Orthophasmavirus coleopterus ICTV
UHM27682.1 RdRp Fushun Phasmavirus 3
QXV86663.1 RdRp Tibet bird virus 1
QMP82227.1 RdRp Coleopteran phasma-related virus OKIAV243
QMP82228.1 RdRp Blattodean phasma-related virus OKIAV239
YP 009304995.1 RdRp Orthophasmavirus wuchangense ICTV
YP 009666959.1 RdRp Seattle Prectang virus
QID77675.1 RdRp Pink bollworm virus 2
YP 009329887.1 RdRp Orthophasmavirus odonatus ICTV
YP 009329871.1 RdRp Orthophasmavirus hubeiense ICTV
YP 010840616.1 RdRp Orthophasmavirus philoctetis ICTV
YP 009666981.1 RdRp Orthophasmavirus gandaense ICTV
YP 010840617.1 RdRp Orthophasmavirus chrysis ICTV
UPT53726.1 RdRp Bactrocera latifrons orthophasmavirus
APG79294.1 RdRp Hubei diptera virus 6
QIJ70043.1 RdRp Peralta bunya-like virus
YP 009553313.1 RdRp Yongsan bunyavirus 1
QIJ56910.1 RdRp Cicadellivirus scaphoidei ICTV
AKN56908.1 RdRp Feravirus ferakinum ICTV
YP 010840748.1 RdRp Guagua virus
YP 010840047.1 RdRp Feravirus neuropterus ICTV
YP 010840003.1 RdRp Feravirus hemipterus ICTV
YP 010085075.1 RdRp Sawastrivirus sanxiaense ICTV
YP 010840610.1 RdRp Hymovirus chrysis ICTV
YP 010840603.1 RdRp Hymovirus chrysurae ICTV
AWA82264.1 RdRp Notori virus
YP 009270651.1 RdRp Wuhivirus insecti ICTV
UZM08148.1 RdRp Aphid bunyavirus 1
YP 009551602.1 RdRp Saesbyeol virus
UHR49905.1 RdRp Sanya sesamia inferens phasmavirus 1
UHK03329.1 RdRp Hangzhou frankliniella intonsa phasmavirus 1
UHK03311.1 RdRp Hangzhou phasmavirus 1
Ab_virus2_seg1_Adaliaphasmavirus germanense_8239nt
YP 009507846.1 RdRp Jonvirus eboris ICTV
ULE62088.1 RdRp Mikado virus
Phenuiviridae
YP 010839720.1 RdRp Bandavirus albatrossense ICTV
YP 008719916.1 RdRp Bandavirus razdanense ICTV
Tree scale: 1
Genera
Orthophasmavirus
Cicadellivirus
Feravirus
Sawastrivirus
Hymovirus
Wuhivirus
Jonvirus
Unclassified
Outgroup

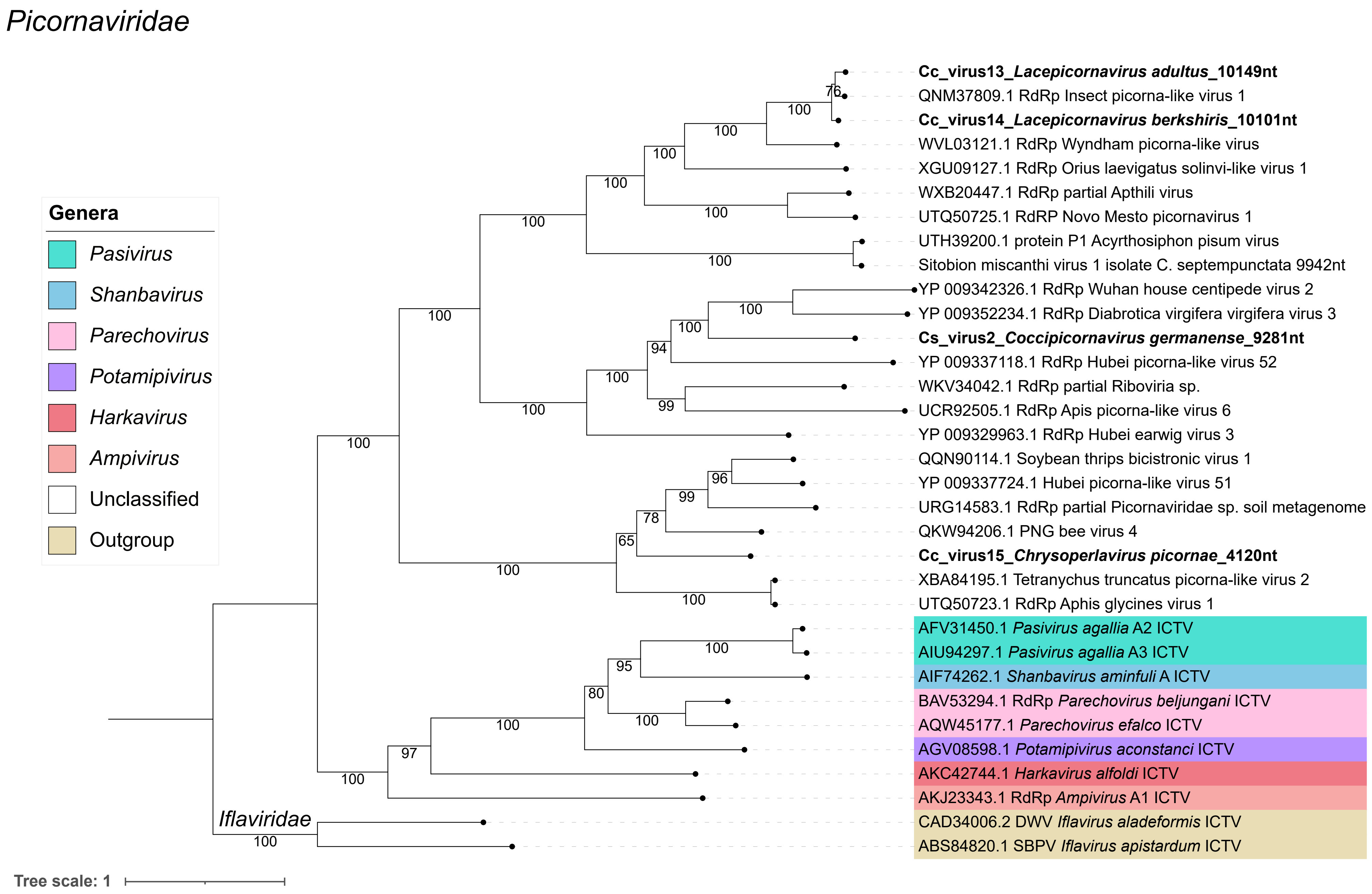
Picornaviridae
Genera
Pasivirus
Shanbavirus
Parechovirus
Potamipivirus
Harkavirus
Ampivirus
Unclassified
Outgroup
Cc_virus13_Lacepicornavirus adultus_10149nt
QNM37809.1 RdRp Insect picorna-like virus 1
Cc_virus14_Lacepicornavirus berkshiris_10101nt
WVL03121.1 RdRp Wyndham picorna-like virus
XGU09127.1 RdRp Orius laevigatus solinvi-like virus 1
WXB20447.1 RdRp partial Apthili virus
UTQ50725.1 RdRP Novo Mesto picornavirus 1
UTH39200.1 protein P1 Acyrthosiphon pisum virus
Sitobion miscanthi virus 1 isolate C. septempunctata 9942nt
YP 009342326.1 RdRp Wuhan house centipede virus 2
YP 009352234.1 RdRp Diabrotica virgifera virgifera virus 3
Cs_virus2_Coccipicornavirus germanense_9281nt
YP 009337118.1 RdRp Hubei picorna-like virus 52
WKV34042.1 RdRp partial Riboviria sp.
UCR92505.1 RdRp Apis picorna-like virus 6
YP 009329963.1 RdRp Hubei earwig virus 3
QQN90114.1 Soybean thrips bicistronic virus 1
YP 009337724.1 Hubei picorna-like virus 51
URG14583.1 RdRp partial Picornaviridae sp. soil metagenome
QKW94206.1 PNG bee virus 4
Cc_virus15_Chrysoperlavirus picornae_4120nt
XBA84195.1 Tetranychus truncatus picorna-like virus 2
UTQ50723.1 RdRp Aphis glycines virus 1
AFV31450.1 Pasivirus agallia A2 ICTV
AIU94297.1 Pasivirus agallia A3 ICTV
AIF74262.1 Shanbavirus aminfuli A ICTV
BAV53294.1 RdRp Parechovirus beljungani ICTV
AQW45177.1 Parechovirus efalco ICTV
AGV08598.1 Potamipivirus aconstanci ICTV
AKC42744.1 Harkavirus alfoldi ICTV
AKJ23343.1 RdRp Ampivirus A1 ICTV
CAD34006.2 DWV Iflavirus aladeformis ICTV
ABS84820.1 SBPV Iflavirus apistardum ICTV
Iflaviridae
76
100
100
100
100
100
100
100
100
100
100
100
94
99
100
100
96
99
78
65
100
100
100
100
100
95
80
100
100
97
100
100
Tree scale: 1

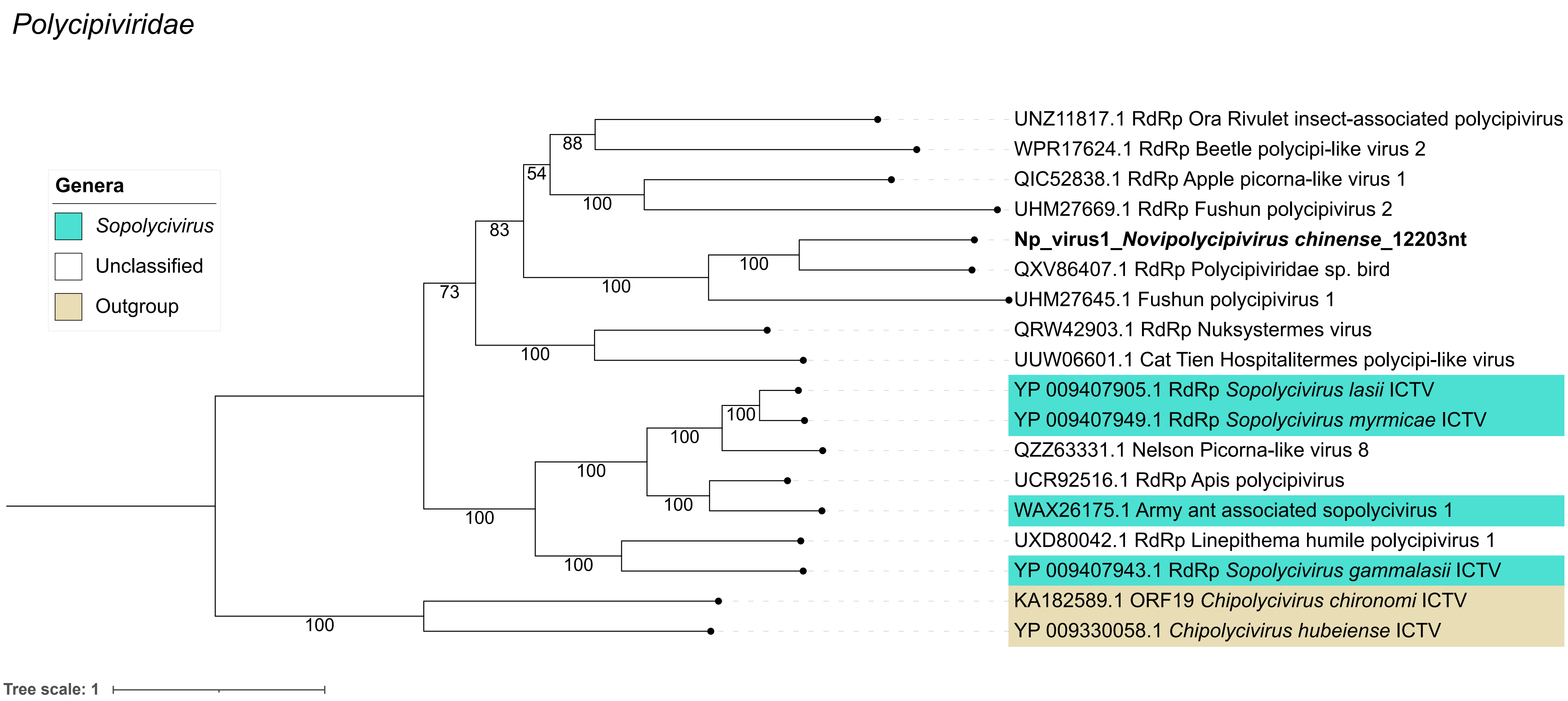

Polycipiviridae
Genera
Sopolycivirus
Unclassified
Outgroup
UNZ11817.1 RdRp Ora Rivulet insect-associated polycipivirus
WPR17624.1 RdRp Beetle polycipi-like virus 2
QIC52838.1 RdRp Apple picorna-like virus 1
UHM27669.1 RdRp Fushun polycipivirus 2
Np_virus1_Novipolycipivirus chinense_12203nt
QXV86407.1 RdRp Polycipiviridae sp. bird
UHM27645.1 Fushun polycipivirus 1
QRW42903.1 RdRp Nuksystermes virus
UUW06601.1 Cat Tien Hospitalitermes polycipi-like virus
YP 009407905.1 RdRp Sopolycivirus lasii ICTV
YP 009407949.1 RdRp Sopolycivirus myrmicae ICTV
QZZ63331.1 Nelson Picorna-like virus 8
UCR92516.1 RdRp Apis polycipivirus
WAX26175.1 Army ant associated sopolycivirus 1
UXD80042.1 RdRp Linepithema humile polycipivirus 1
YP 009407943.1 RdRp Sopolycivirus gammalasii ICTV
KA182589.1 ORF19 Chipolycivirus chironomi ICTV
YP 009330058.1 Chipolycivirus hubeiense ICTV
88
54
100
83
100
100
73
100
100
100
100
100
100
100
100
Tree scale: 1

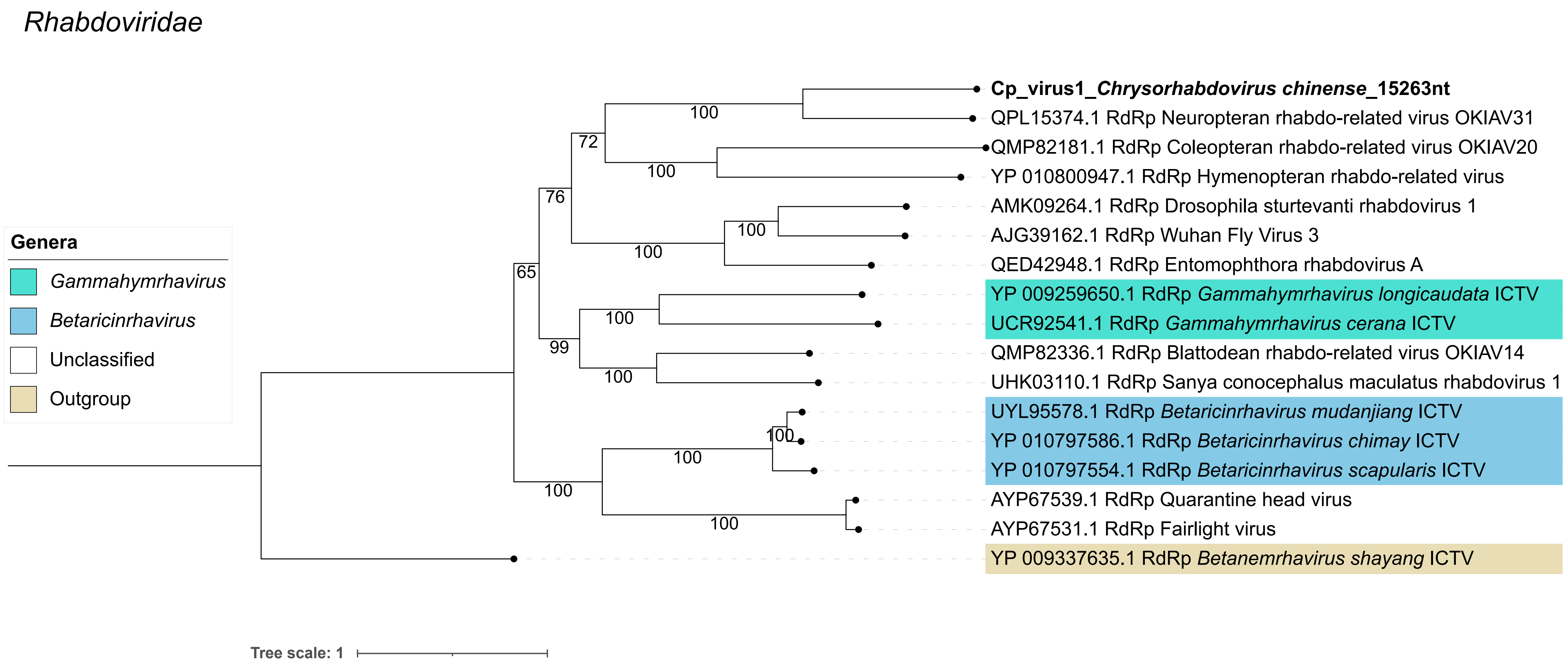

Rhabdoviridae
Genera
Gammahymrhavirus
Betaricinrhavirus
Unclassified
Outgroup
Cp_virus1_Chrysorhabdovirus chinense_15263nt
QPL15374.1 RdRp Neuropteran rhabdo-related virus OKIAV31
QMP82181.1 RdRp Coleopteran rhabdo-related virus OKIAV20
YP 010800947.1 RdRp Hymenopteran rhabdo-related virus
AMK09264.1 RdRp Drosophila sturtevanti rhabdovirus 1
AJG39162.1 RdRp Wuhan Fly Virus 3
QED42948.1 RdRp Entomophthora rhabdovirus A
YP 009259650.1 RdRp Gammahymrhavirus longicaudata ICTV
UCR92541.1 RdRp Gammahymrhavirus cerana ICTV
QMP82336.1 RdRp Blattodean rhabdo-related virus OKIAV14
UHK03110.1 RdRp Sanya conocephalus maculatus rhabdovirus 1
UYL95578.1 RdRp Betaricinrhavirus mudanjiang ICTV
YP 010797586.1 RdRp Betaricinrhavirus chimay ICTV
YP 010797554.1 RdRp Betaricinrhavirus scapularis ICTV
AYP67539.1 RdRp Quarantine head virus
AYP67531.1 RdRp Fairlight virus
YP 009337635.1 RdRp Betanemrhavirus shayang ICTV
100
72
100
76
100
100
65
100
99
100
100
100
100
100
Tree scale: 1

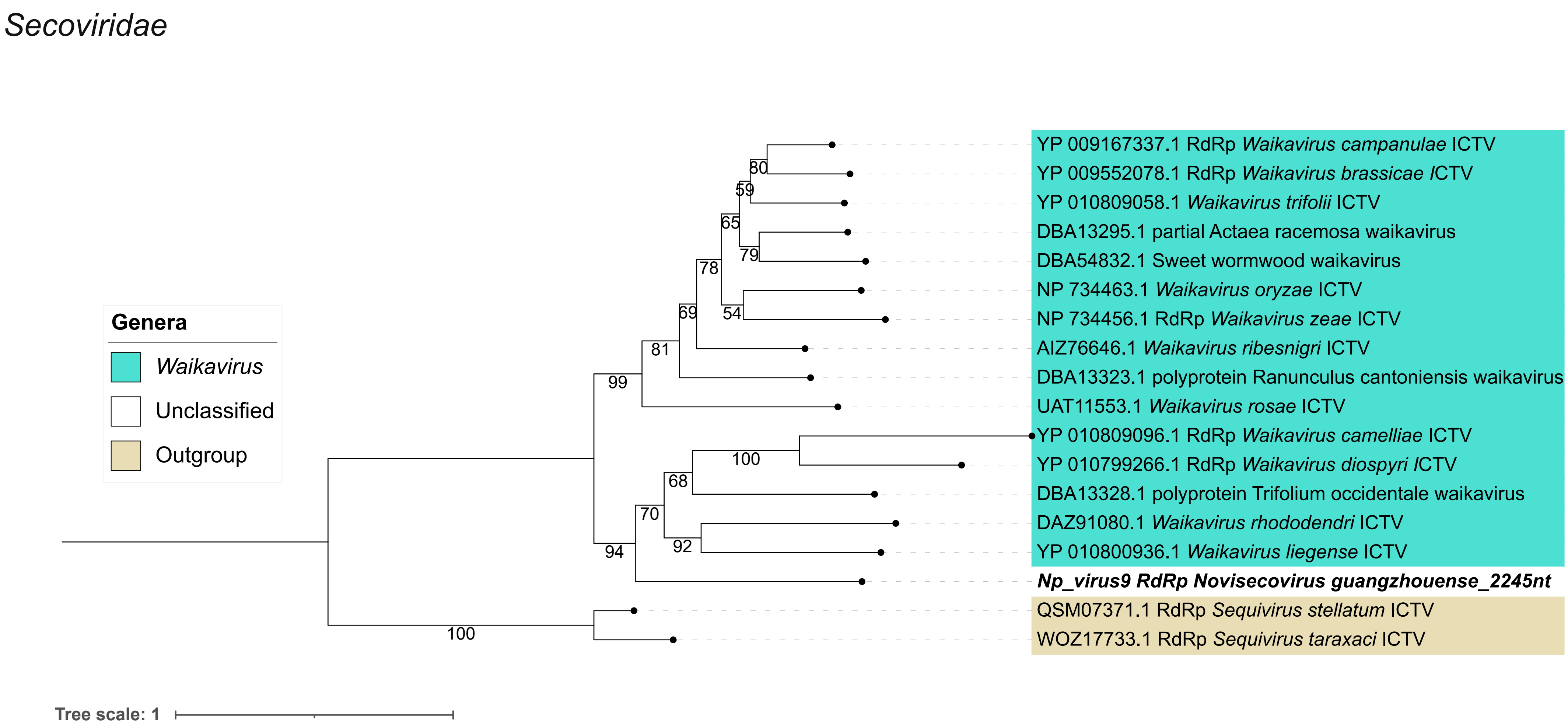
Secoviridae
Genera
Waikavirus
Unclassified
Outgroup
YP 009167337.1 RdRp Waikavirus campanulae ICTV
YP 009552078.1 RdRp Waikavirus brassicae ICTV
YP 010809058.1 Waikavirus trifolii ICTV
DBA13295.1 partial Actaea racemosa waikavirus
DBA54832.1 Sweet wormwood waikavirus
NP 734463.1 Waikavirus oryzae ICTV
NP 734456.1 RdRp Waikavirus zeae ICTV
AIZ76646.1 Waikavirus ribesnigri ICTV
DBA13323.1 polyprotein Ranunculus cantoniensis waikavirus
UAT11553.1 Waikavirus rosae ICTV
YP 010809096.1 RdRp Waikavirus camelliae ICTV
YP 010799266.1 RdRp Waikavirus diospyri ICTV
DBA13328.1 polyprotein Trifolium occidentale waikavirus
DAZ91080.1 Waikavirus rhododendri ICTV
YP 010800936.1 Waikavirus liegense ICTV
Np_virus9 RdRp Novisecovirus guangzhouense_2245nt
QSM07371.1 RdRp Sequivirus stellatum ICTV
WOZ17733.1 RdRp Sequivirus taraxaci ICTV
80
59
65
79
78
54
69
81
99
100
68
70
92
94
100
Tree scale: 1

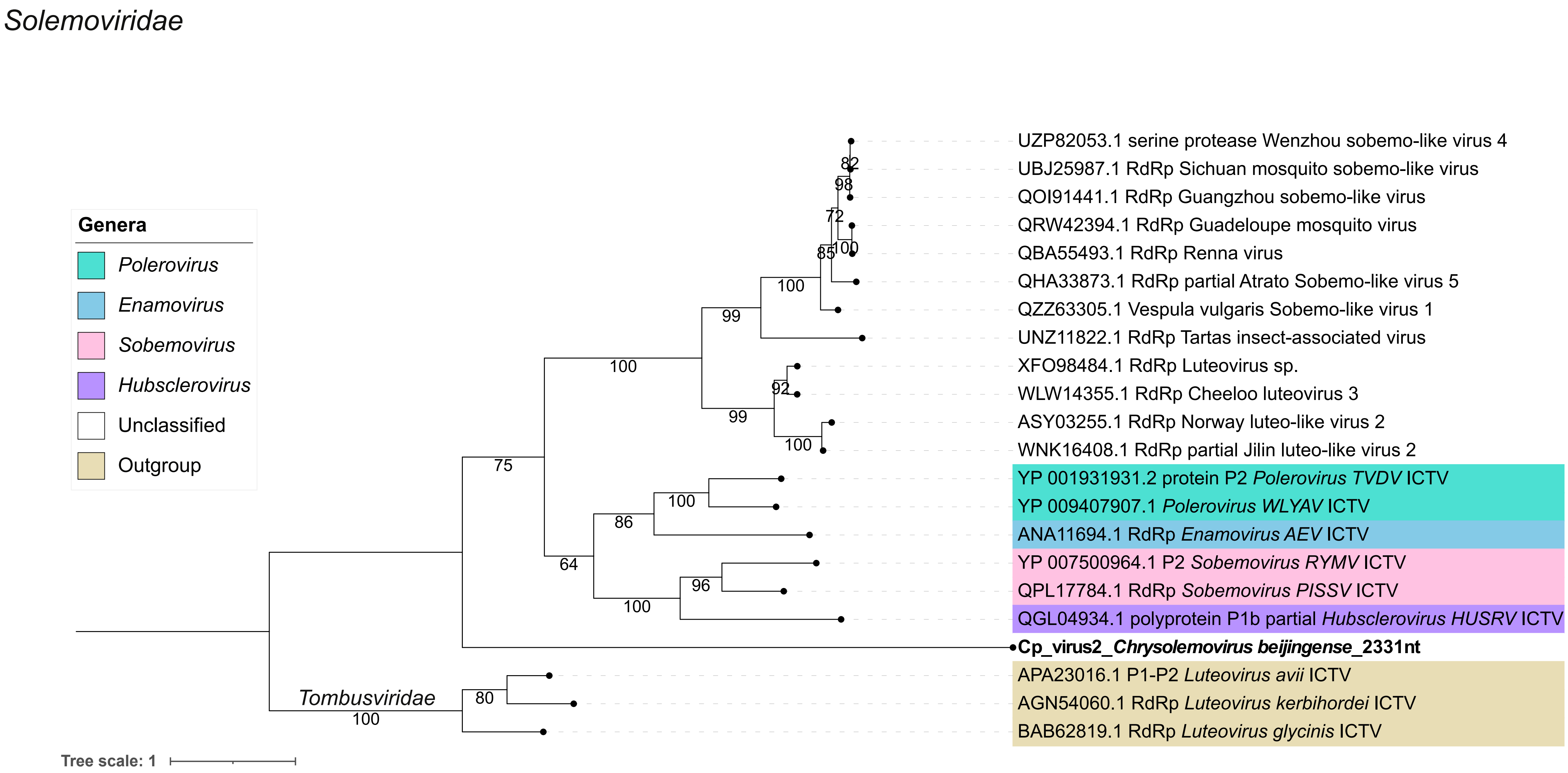
Solemoviridae
Genera
Polerovirus
Enamovirus
Sobemovirus
Hubsclerovirus
Unclassified
Outgroup
UZP82053.1 serine protease Wenzhou sobemo-like virus 4
UBJ25987.1 RdRp Sichuan mosquito sobemo-like virus
QOI91441.1 RdRp Guangzhou sobemo-like virus
QRW42394.1 RdRp Guadeloupe mosquito virus
QBA55493.1 RdRp Renna virus
QHA33873.1 RdRp partial Atrato Sobemo-like virus 5
QZZ63305.1 Vespula vulgaris Sobemo-like virus 1
UNZ11822.1 RdRp Tartas insect-associated virus
XFO98484.1 RdRp Luteovirus sp.
WLW14355.1 RdRp Cheeloo luteovirus 3
ASY03255.1 RdRp Norway luteo-like virus 2
WNK16408.1 RdRp partial Jilin luteo-like virus 2
YP 001931931.2 protein P2 Polerovirus TVDV ICTV
YP 009407907.1 Polerovirus WLYAV ICTV
ANA11694.1 RdRp Enamovirus AEV ICTV
YP 007500964.1 P2 Sobemovirus RYMV ICTV
QPL17784.1 RdRp Sobemovirus PISSV ICTV
QGL04934.1 polyprotein P1b partial Hubsclerovirus HUSRV ICTV
Cp_virus2_Chrysolemovirus beijingense_2331nt
APA23016.1 P1-P2 Luteovirus avii ICTV
AGN54060.1 RdRp Luteovirus kerbihordei ICTV
BAB62819.1 RdRp Luteovirus glycinis ICTV
Tombusviridae
82
98
72
85
100
100
99
100
92
99
100
75
100
86
64
96
100
80
100
Tree scale: 1

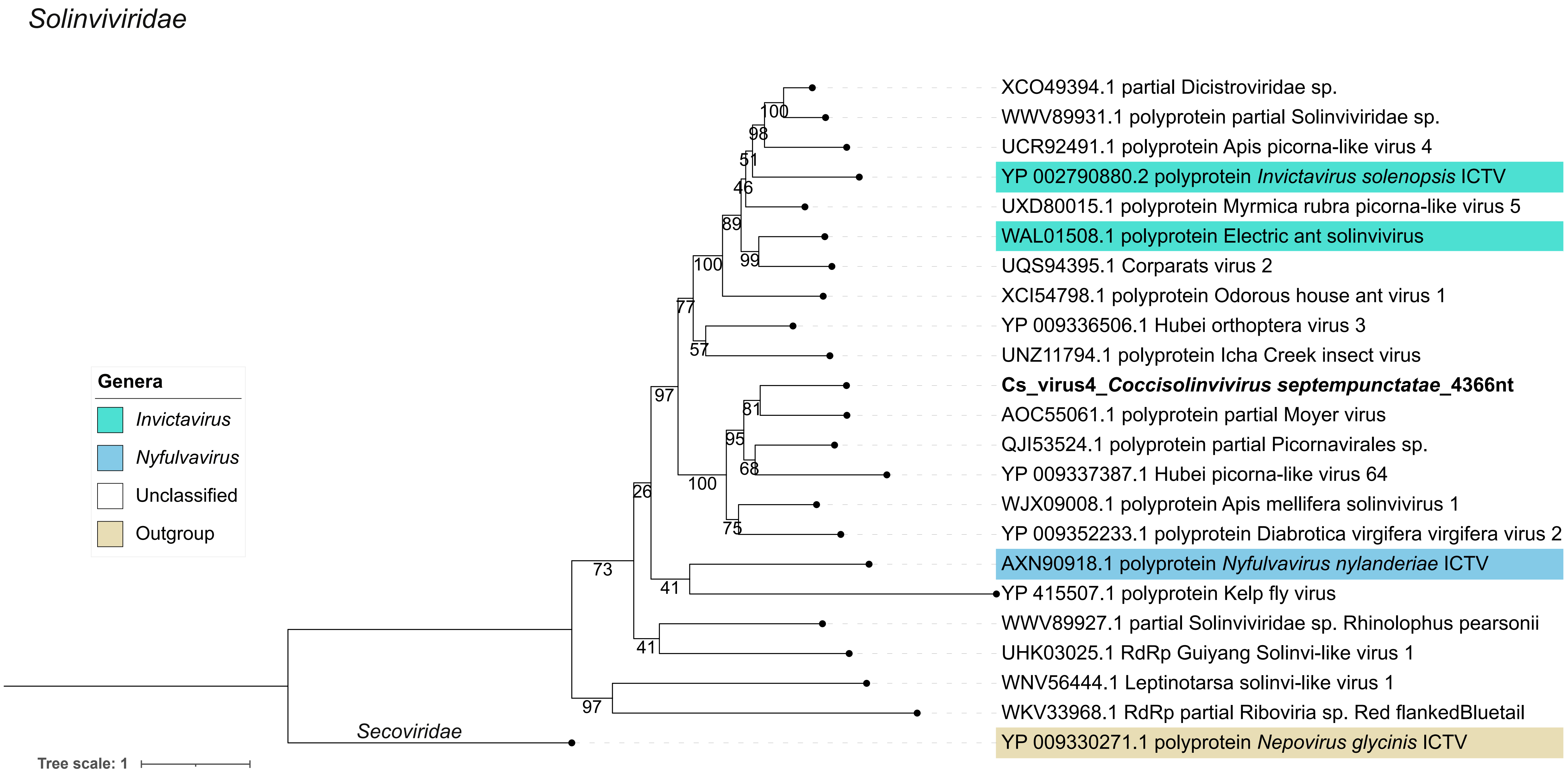

Solinviviridae
XCO49394.1 partial Dicistroviridae sp.
WWV89931.1 polyprotein partial Solinviviridae sp.
UCR92491.1 polyprotein Apis picorna-like virus 4
YP 002790880.2 polyprotein Invictavirus solenopsis ICTV
UXD80015.1 polyprotein Myrmica rubra picorna-like virus 5
WAL01508.1 polyprotein Electric ant solinvivirus
UQS94395.1 Corparats virus 2
XCI54798.1 polyprotein Odorous house ant virus 1
YP 009336506.1 Hubei orthoptera virus 3
UNZ11794.1 polyprotein Icha Creek insect virus
Cs_virus4_Coccisolinvivirus septempunctatae_4366nt
AOC55061.1 polyprotein partial Moyer virus
QJI53524.1 polyprotein partial Picornavirales sp.
YP 009337387.1 Hubei picorna-like virus 64
WJX09008.1 polyprotein Apis mellifera solinvivirus 1
YP 009352233.1 polyprotein Diabrotica virgifera virgifera virus 2
AXN90918.1 polyprotein Nyfulvavirus nylanderiae ICTV
YP 415507.1 polyprotein Kelp fly virus
WWV89927.1 partial Solinviviridae sp. Rhinolophus pearsonii
UHK03025.1 RdRp Guiyang Solinvi-like virus 1
WNV56444.1 Leptinotarsa solinvi-like virus 1
WKV33968.1 RdRp partial Riboviria sp. Red flankedBluetail
YP 009330271.1 polyprotein Nepovirus glycinis ICTV
Secoviridae
100
98
51
46
89
99
100
77
57
97
81
95
68
100
75
26
73
41
41
97
Genera
Invictavirus
Nyfulvavirus
Unclassified
Outgroup
Tree scale: 1

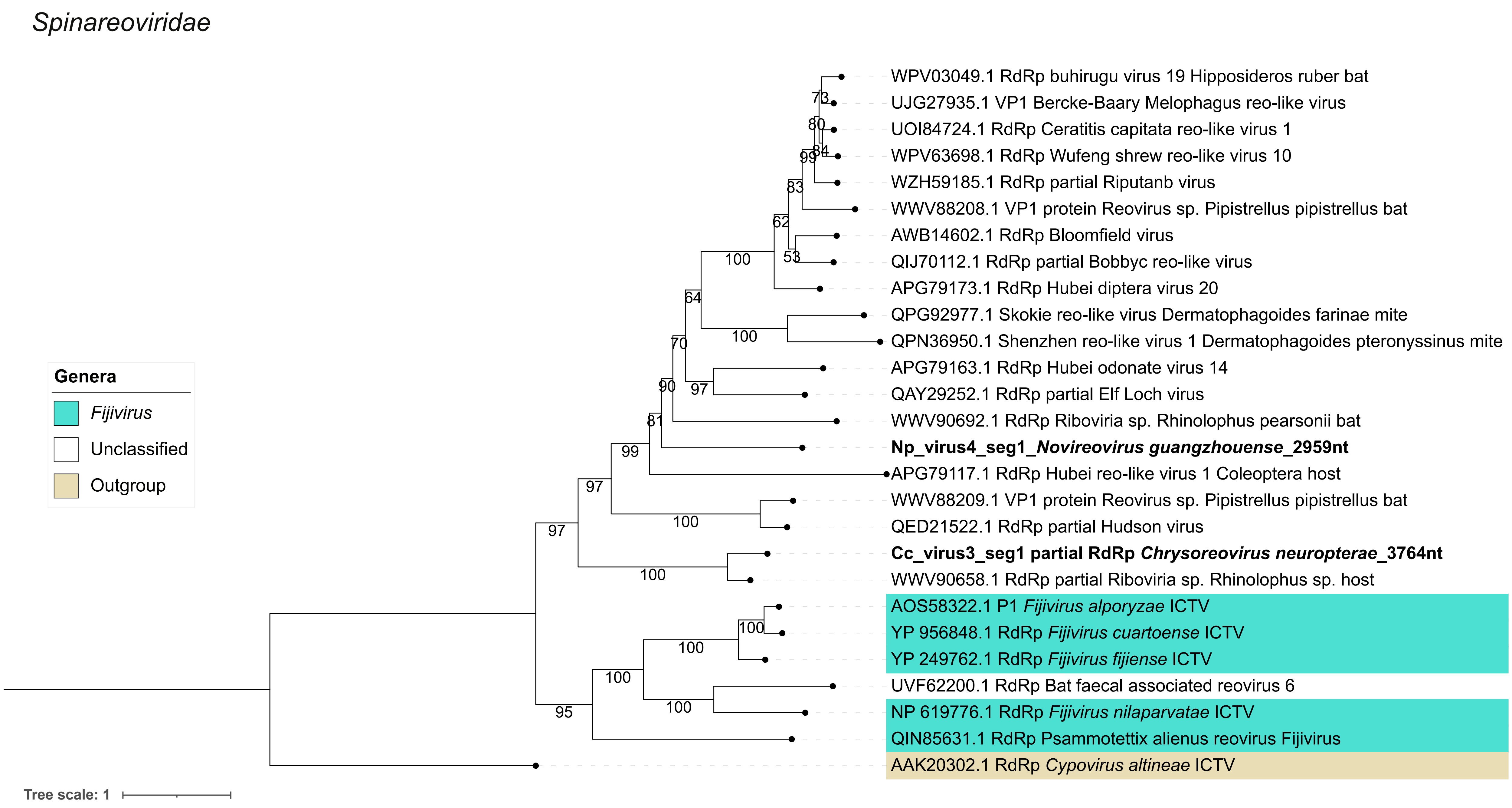
Spinareoviridae
Genera
Fijivirus
Unclassified
Outgroup
WPV03049.1 RdRp buhirugu virus 19 Hipposideros ruber bat
UJG27935.1 VP1 Bercke-Baary Melophagus reo-like virus
UOI84724.1 RdRp Ceratitis capitata reo-like virus 1
WPV63698.1 RdRp Wufeng shrew reo-like virus 10
WZH59185.1 RdRp partial Riputanb virus
WWV88208.1 VP1 protein Reovirus sp. Pipistrellus pipistrellus bat
AWB14602.1 RdRp Bloomfield virus
QIJ70112.1 RdRp partial Bobbyc reo-like virus
APG79173.1 RdRp Hubei diptera virus 20
QPG92977.1 Skokie reo-like virus Dermatophagoides farinae mite
QPN36950.1 Shenzhen reo-like virus 1 Dermatophagoides pteronyssinus mite
APG79163.1 RdRp Hubei odonate virus 14
QAY29252.1 RdRp partial Elf Loch virus
WWV90692.1 RdRp Riboviria sp. Rhinolophus pearsonii bat
Np_virus4_seg1_Novireovirus guangzhouense_2959nt
APG79117.1 RdRp Hubei reo-like virus 1 Coleoptera host
WWV88209.1 VP1 protein Reovirus sp. Pipistrellus pipistrellus bat
QED21522.1 RdRp partial Hudson virus
Cc_virus3_seg1 partial RdRp Chrysoreovirus neuropterae_3764nt
WWV90658.1 RdRp partial Riboviria sp. Rhinolophus sp. host
AOS58322.1 P1 Fijivirus alporyzae ICTV
YP 956848.1 RdRp Fijivirus cuartoense ICTV
YP 249762.1 RdRp Fijivirus fijiense ICTV
UVF62200.1 RdRp Bat faecal associated reovirus 6
NP 619776.1 RdRp Fijivirus nilaparvatae ICTV
QIN85631.1 RdRp Psammotettix alienus reovirus Fijivirus
AAK20302.1 RdRp Cypovirus altineae ICTV
Tree scale: 1

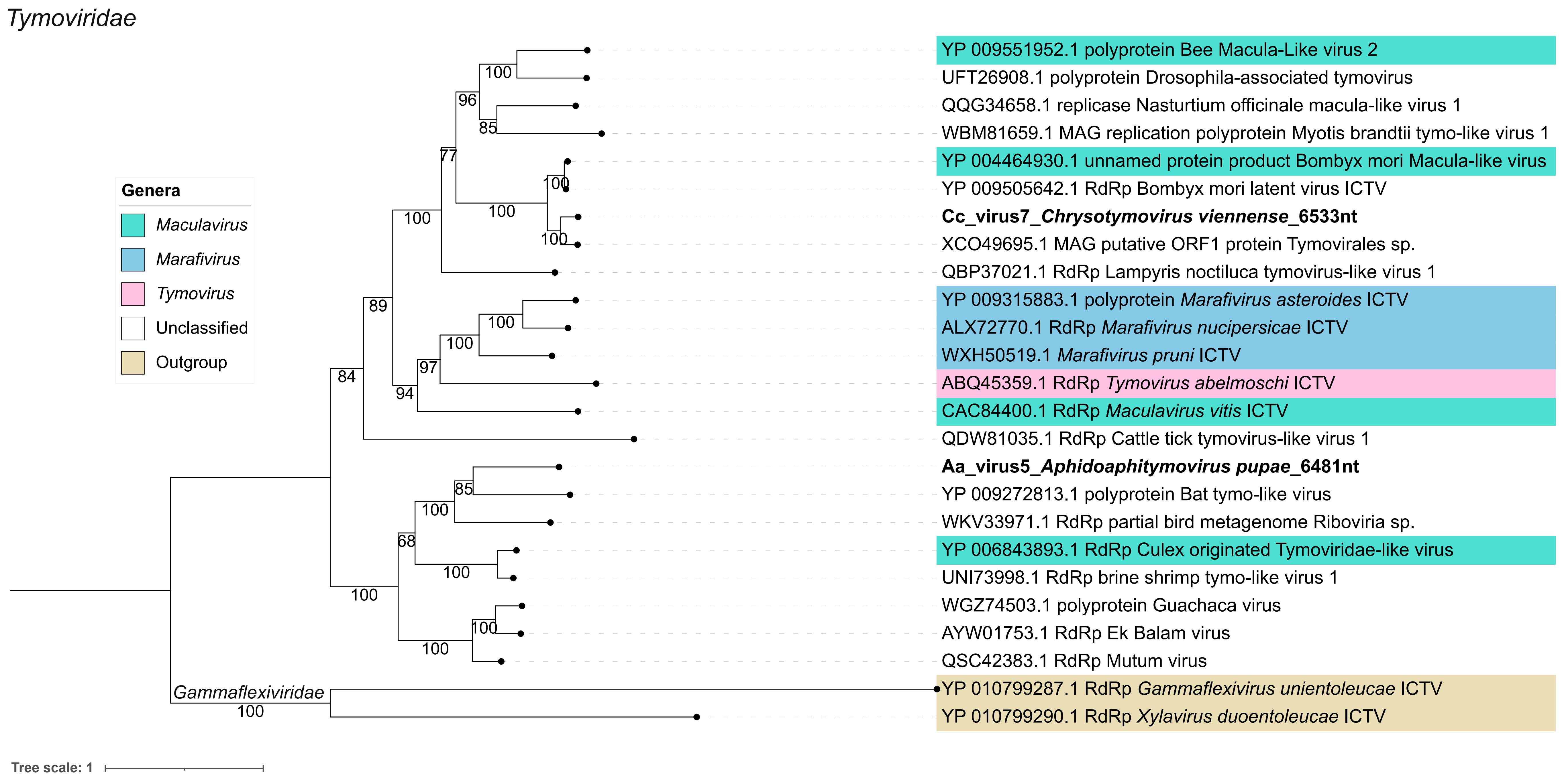
Tymoviridae
Genera
Maculavirus
Marafivirus
Tymovirus
Unclassified
Outgroup
YP 009551952.1 polyprotein Bee Macula-Like virus 2
UFT26908.1 polyprotein Drosophila-associated tymovirus
QQG34658.1 replicase Nasturtium officinale macula-like virus 1
WBM81659.1 MAG replication polyprotein Myotis brandtii tymo-like virus 1
YP 004464930.1 unnamed protein product Bombyx mori Macula-like virus
YP 009505642.1 RdRp Bombyx mori latent virus ICTV
Cc_virus7_Chrysotymovirus viennense_6533nt
XCO49695.1 MAG putative ORF1 protein Tymovirales sp.
QBP37021.1 RdRp Lampyris noctiluca tymovirus-like virus 1
YP 009315883.1 polyprotein Marafivirus asteroides ICTV
ALX72770.1 RdRp Marafivirus nucipersicae ICTV
WXH50519.1 Marafivirus pruni ICTV
ABQ45359.1 RdRp Tymovirus abelmoschi ICTV
CAC84400.1 RdRp Maculavirus vitis ICTV
QDW81035.1 RdRp Cattle tick tymovirus-like virus 1
Aa_virus5_Aphidoaphitymovirus pupae_6481nt
YP 009272813.1 polyprotein Bat tymo-like virus
WKV33971.1 RdRp partial bird metagenome Riboviria sp.
YP 006843893.1 RdRp Culex originated Tymoviridae-like virus
UNI73998.1 RdRp brine shrimp tymo-like virus 1
WGZ74503.1 polyprotein Guachaca virus
AYW01753.1 RdRp Ek Balam virus
QSC42383.1 RdRp Mutum virus
YP 010799287.1 RdRp Gammaflexivirus unientoleucae ICTV
YP 010799290.1 RdRp Xylavirus duoentoleucae ICTV
Gammaflexiviridae
Tree scale: 1
